# Successive passaging of a plant-associated microbiome reveals robust habitat and host genotype-dependent selection

**DOI:** 10.1101/627794

**Authors:** Norma M. Morella, Francis Cheng-Hsuan Weng, Pierre M. Joubert, C. Jessica E. Metcalf, Steven Lindow, Britt Koskella

## Abstract

There is increasing interest in the plant microbiome as it relates to both plant health and agricultural sustainability. One key unanswered question is whether we can select for a plant microbiome that is robust after colonization of target hosts. We used a successive passaging experiment to address this question by selecting upon the tomato phyllosphere microbiome. Beginning with a diverse microbial community generated from field-grown tomato plants, we inoculated replicate plants across five plant genotypes for four eight-week long passages, sequencing the microbial community at each passage. We observed consistent shifts in both the bacterial (16S amplicon sequencing) and fungal (ITS amplicon sequencing) communities across replicate lines over time, as well as a general loss of diversity over the course of the experiment suggesting that much of the naturally observed microbial community in the phyllosphere is likely transient or poorly adapted. We found that both host genotype and environment shape microbial composition, but the relative importance of genotype declines through time. Furthermore, using a community coalescence experiment, we found that the bacterial community from the end of the experiment was robust to invasion by the starting bacterial community. These results highlight that selecting for a stable microbiome that is well adapted to a particular host environment is indeed possible, emphasizing the great potential of this approach in agriculture and beyond.

**Significance Statement:** There is great interest in selecting for host-associated microbiomes that confer particular functions to their host, and yet it remains unknown whether selection for a robust and stable microbiome is possible. Here, we use a microbiome passaging approach to measure the impact of host-mediated selection on the tomato phyllosphere (above ground) microbiome. We find robust community selection across replicate lines that is shaped by plant host genotype in early passages, but changes in a genotype-independent manner in later passages. Work such as ours is crucial to understanding the general principles governing microbiome assembly and adaptation, and is widely applicable to both sustainable agriculture and microbiome-related medicine.

## Introduction

The study of microbiomes (diverse microbial communities and their collective genomes) spans both basic and applied research in human health, agriculture, and environmental change. As our understanding of the ability of the microbiome to influence host health and shape host traits deepens, there is increasing interest in selecting and/or designing microbiomes for specific traits or functions. Such trait-based selection of microbiomes has the potential to shape the future of agriculture and medicine [1][2]. In agriculture, below-ground microbiota have already proven capable of shifting the flowering time of plant hosts [3], enhancing drought resistance [4, 5], and even altering above-ground herbivory [6]. However, long-term, repeatable success of future efforts will rely on a fundamental understanding of the assembly of, selection within, and co-evolution among microbiota within these communities. One of the challenges facing successful, rational microbiome manipulation and assembly is disentangling the forces naturally shaping the communities, including both host characteristics and microbial immigration on community stability. For example, in both humans and plants, there is conflicting evidence as to the relative importance of the environment versus host genotype in shaping the microbiome [7–15], and dispersal has been shown to override host genetics in an experimental zebra fish system [16].

One powerful but under-utilized approach to understand and experimentally control for the factors shaping microbiome composition and diversity is experimental evolution. Measuring changes of populations or communities over time under controlled settings in response to a known selection pressure has proved a powerful force in gaining fundamental understanding of both host-pathogen (co)evolution [17] and microbial evolution [18]. Here, we harness an experimental evolution approach in order to study how an entire microbial community can be selected upon in a plant host environment that varies across disease resistance-associated genotypes. We test the fundamental yet relatively untested assumption that a microbiome can be selected to adapt to its host in a robust fashion. To do this, we employ a microbiome passaging approach using the phyllosphere microbiome of tomato (*Solanum*) as a model system to select for a community that is capable of growth in this relatively oligotrophic environment and is resilient to perturbation via competition with a non-‘adapted,’ but more diverse community. The phyllosphere, defined as the aerial surfaces of the plant, is a globally important microbial habitat [19], and can shape important plant traits such as protection against foliar disease [20, 21] and growth [22, 23]. Successful trait-based selection on the phyllosphere could therefore allow for enhancement of plant health, but this critically depends on the ability to select for a well-adapted microbial community that is relatively stable against invasion, particularly in open environments in which dispersal from neighboring hosts or the surrounding environment is inevitable.

We collected a diverse phyllosphere microbiome from tomatoes grown in an agricultural setting and transplanted it onto green-house grown plants using a transplantation method previously shown to be effective for lettuce [24]. We serially passaged this diverse microbiome on each of four cohorts of tomato plants (six lines per cohort) of five different genotypes (pairs of near isogenic *S. lycopersicum* genotypes that differed at known disease resistance loci, as well as a wild tomato accession, *S. pimpinellifolium*) for a total of 30 weeks. On each plant, during each passage, community assembly and dynamics might be driven by neutral processes or reflect positive or negative selection of specific taxa by the plant, the greenhouse environment, and/or the other microbial taxa present. We therefore sought to characterize the relative importance of neutral versus deterministic processes both computationally using a neutral model, and empirically using community coalescence experiments [25] in which communities from different passaged lines are combined together and re-inoculated onto host plants in a common garden experiment. Overall, we were able to measure and characterize the response of the phyllosphere microbiome to selection in the plant host environment under greenhouse conditions, and our findings suggest selection for a stable and well-adapted plant-associated microbiome.

## Results

### Serial passaging experiment

A diverse starting inoculum was collected from field grown, mature tomato plants. This field-microbiome was spray inoculated onto 30 tomato plants of 5 different genotypes, with six replicates each. Two-week old tomato plants were spray-inoculated once per week for five weeks, and then sampled in their entirety ten days after the final inoculation (Figure 1b). The phyllosphere microbiome of each plant was then individually passaged on these genetically distinct hosts over the course of four eight-week long passages; P1, P2, P3, and P4 (Figure 1a; see methods for details). Microbiomes were not pooled across plants within a given plant genotype, resulting in 30 independent selection lines. Control plants were inoculated with an equal volume of either heat killed inoculum (P1) or sterile buffer (subsequent passages) every week. At the end of each passage, bacterial density was measured and normalized to the weight of each plant (Figure 1c), and communities were sequenced using 16S rRNA amplicon sequencing.

**Figure 1.**
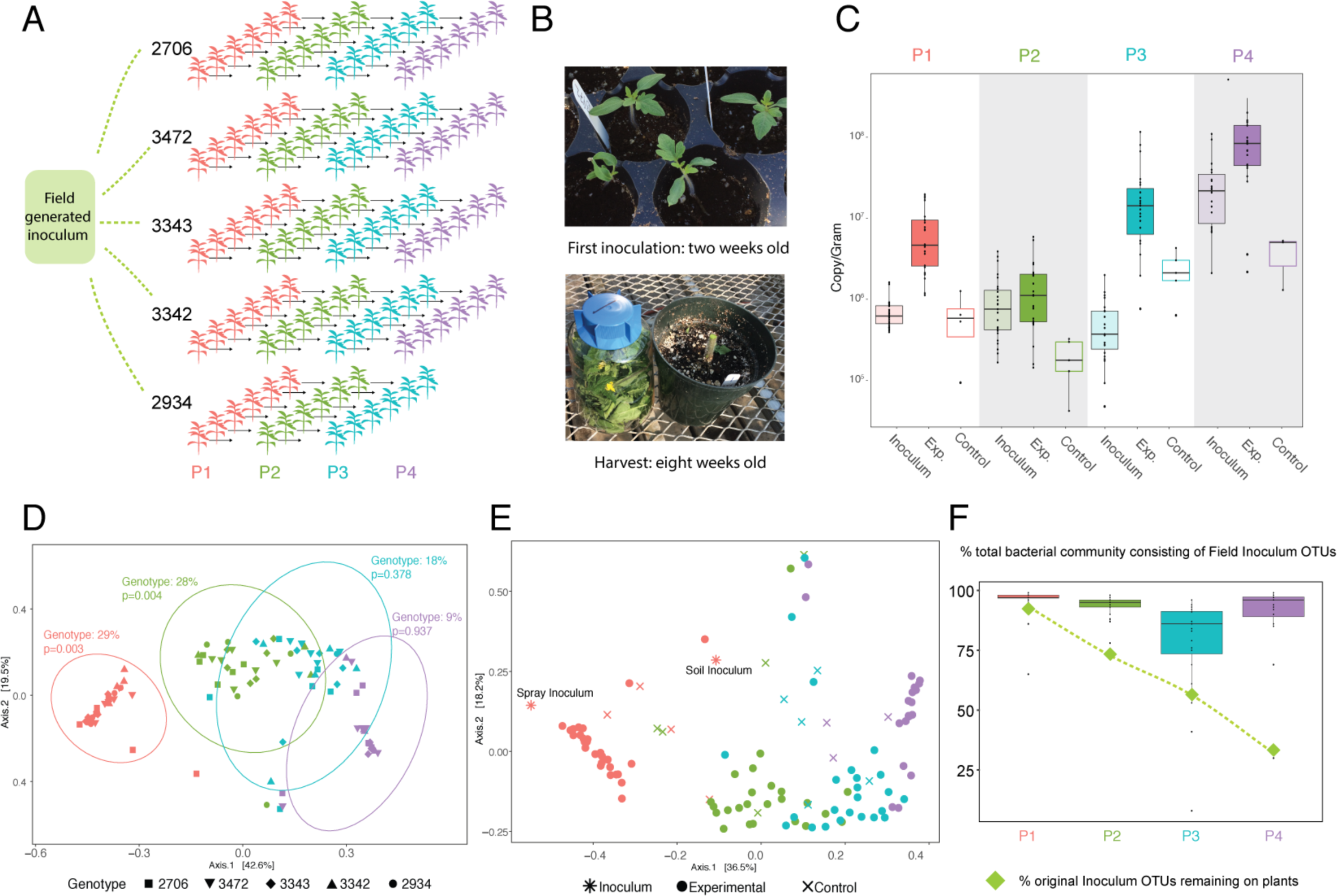
Serial passaging of the phyllosphere microbiome. Experimental design of serial passaging experiment where microbial inoculum from an agricultural tomato field was inoculated onto replicates of five genotypes and passaged for four passages (a). Plants were first inoculated when they were 2-weeks old, and the entire plant was sampled at 8 weeks old (b). Bacterial abundance was measured at the end of each passage from experimental and control plants using ddPCR and normalized to the weight of each plant. Inoculum density was calculated as well (c). PCoA plots of Bray-Curtis distances show a significant effect (determined by a PERMANOVA test) of genotype in P1 and P2 (d) and passage (colors) and sample type (shapes) (e). Ellipses indicate 95% confidence around the clustering. The percent of original inoculum OTUs present at each passage was calculated (green diamonds), and the reads/sample of inoculum OTUs out of total reads was calculated for each plant at every passage and displayed on a box plot (f).

We first measured the impact of host genotype on bacterial community structure (Figure 1d). Using Bray-Curtis dissimilarity measures, we performed permutational multivariate analysis of variance tests using the Vegan’s Adonis function and found that plant genotype explains 29% of dissimilarity between microbiomes in P1 (p=0.003). In P2, plant genotype similarly explains 28% of the variation in bacterial community dissimilarity (p=0.004). However, genotype becomes an insignificant driver of community composition in both P3 (18%, p=0.378) and P4 (9%, p=0.937) and is robust to the removal of the outlying sample in P1 (see supplemental methods).

We also sought to determine if there were more subtle influences of host genotype on the community that were not uncovered through analyzing Bray-Curtis distances alone. From the original inoculum sample, we identified ten Operational Taxonomic Units (OTUs) using linear discriminant analysis effect-size (LEfSe) analysis [26] that were significantly associated with particular genotypes in P1 and P2. We compared their presence/absence at the end of P4 to those OTUs that were not found to be associated with genotype. Interestingly, those OTUs that were significantly associated with particular genotypes at the start of the experiment were significantly more likely to be present at the end of the experiment than those not associated with genotype (Fisher’s exact test, p=0.013).

In addition to genotype effects, we were interested in what other factors were driving our observed change in community composition. We found that the number of passages on tomato plants strongly shaped microbial community diversity. Bray-Curtis distances across all samples uncovered a significant effect of both passage number and sample type (i.e. experimental, control, or inoculum) on bacterial communities (Figure 1e; effect of Passage F_3,_ 114= 27.8895, p= 0.001; Sample Type F_3, 114_= 3.0075, p=0.001). As this was an open system, we next sought to determine if there was a high degree of dispersal amongst plants within the greenhouse by directly comparing the communities of experimental and control plants. At every passage, control and experimental plants are found to host significantly different communities (all p-values <0.04), suggesting minimal effects of dispersal within the greenhouse relative to our inoculations. When inoculum and control samples are removed from analysis, there remains a significant effect of passage number (F_3, 89_= 33.3023p=0.001) and a significant overall effect of plant genotype on community composition (F_4, 89_= 1.9991, p=0.016). When variance is partitioned, passage can explain 51% of dissimilarity, whereas genotype explains only 4%. Replicate lines from accession 2934 were lost after P3 due to a stem rot fungal pathogen present in the original inoculum that seemingly only infected this genotype. However, the observed overall genotype effect was not driven by this accession, as there remains a significant effect of genotype after its removal (F_3, 79_= 1.9723, p= 0.034), and passage number remains highly significant (F_3, 79_= 31.9804, p= 0.001).

To better understand how the original, diverse, field inoculum changed over four passages on plants in the greenhouse, we calculated the percentage of OTUs in the original inoculum that were detectable over the course of the experiment (Figure 1f, green diamonds). At the end of P1, 92% of the field inoculum OTUs were still present on the plants, but by P4, this was reduced to 29%. We then calculated if the decrease in original community member diversity was the result of replacement by non-inoculum taxa (i.e. those that colonized plants over the course of the experiment). In this case, we observed that the proportion of sequencing reads (divided by total reads) representing the original inoculum OTUs remains above 78% (Figure 1f, box plots). This suggests that a relatively small percentage of the community was made up of OTUs that colonized plants from the greenhouse environment. Of note, some OTUs considered “non-inoculum” were likely present in the initial inoculum, but in too low of abundance to detect. In particular, there were 27 OTUs with reads in the spray inoculum sample in the non-rarefied dataset, but this was number was reduced to zero after data rarefaction. To account for the impact of the small percentage of arriving species on community composition, we re-analyzed the dataset using only those OTUs that were observed to be present in the initial inoculum (Supplemental Figure S1a). In this case, passage number remains a significant driver of community dissimilarity (F_3 89_= 37.6813, p=0.001), as does genotype (F_4, 89_= 2.0393, p=0.015).

We next measured changes in community diversity over the course of passaging and across lines. We found a significant decrease in both OTU richness and alpha diversity over time across all plant genotypes (Figure 2a and b), including when only original spray inoculum OTUs are considered (Supplemental Figure S1b). Importantly, this drop in diversity from the start of the experiment does not correspond to a decrease in overall bacterial abundance on plants (Figure 1b). Note that our measures of bacterial growth likely largely overestimate the starting densities and do not account for population turnover (as a result of cell death and replacement within a passage), and are therefore highly conservative. In P1, we also estimated fold change of bacterial abundance on control plants that were sprayed with heat-killed inoculum, and found an average change of 0.76, which is significantly lower than the averaged 11-fold change for experimental plants which received live inoculum (Welch’s Two sample T-Test, p<0.0001). Finally, although passaging was performed in a control temperature greenhouse, outside high and low temperatures and humidity all varied significantly across passages (Supplemental Figure 2; ANOVA P<0.001 for all measures), which may have impacted the observed differences in both abundance and growth across passages.

**Figure 2.**
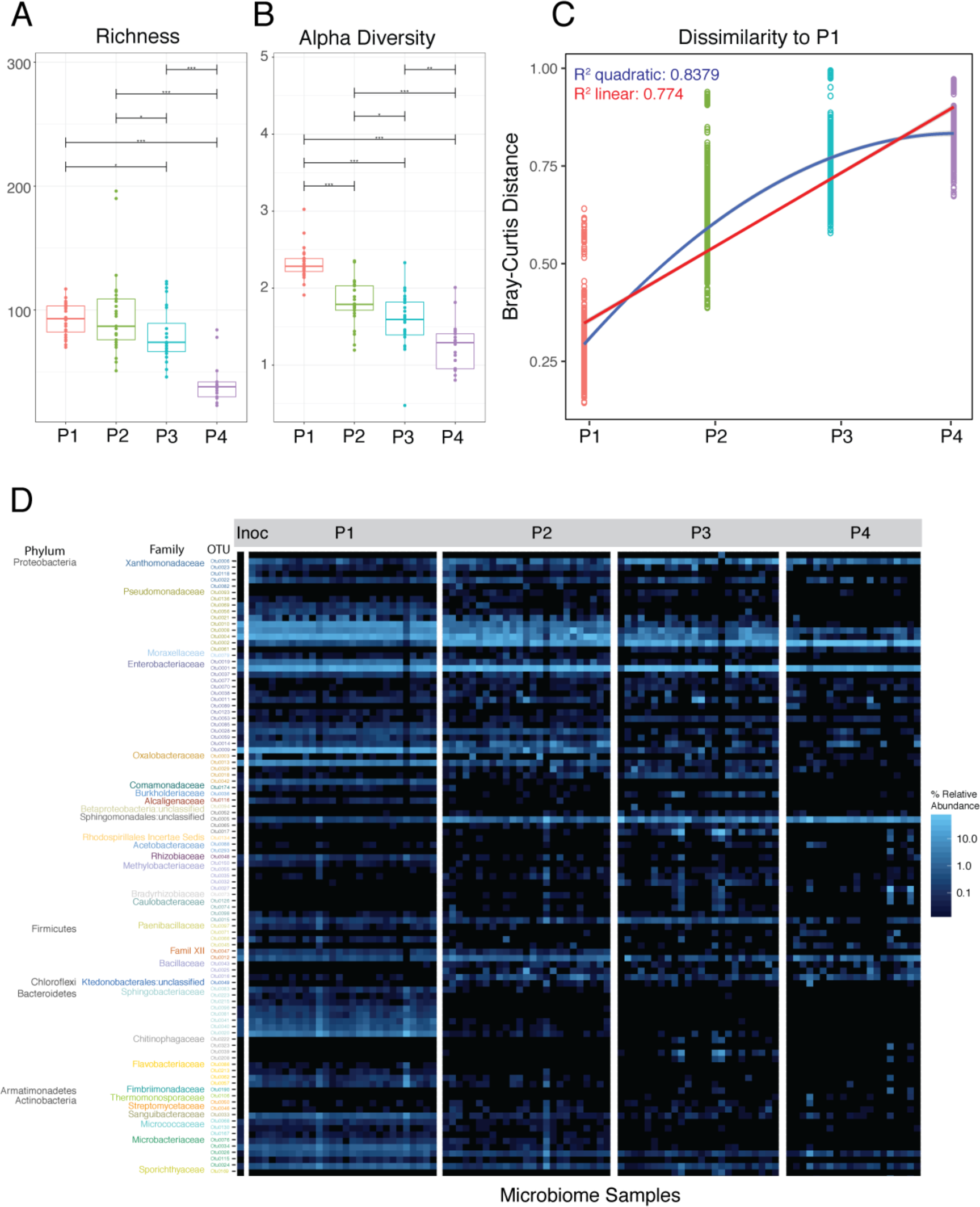
Changes in diversity and composition from P1 to P4. Plots of richness (a) and Shannon’s alpha diversity (b) at each passage show a significant decrease over time. Bray Curtis distances between microbiomes in P1 were compared to those in P1, P2, P3, and P4, and linear and quadratic models were fit to the data (c). A heat map showing relative abundance of the top 100 OTUs illustrates the changing community composition at multiple taxonomic levels (d). Full taxonomy of OTUs is found in Supplemental Table 1. Significance values of pairwise comparisons in (a) and (b) are illustrated on the graph * p≤0.05; ** p≤0.01; *** p≤0.001; ****p≤0.0001.

With the knowledge that communities were drastically changing over time, we sought to determine if the rate at which the communities were changing was consistent. To do this, we calculated Bray-Curtis distances of microbiomes in each passage to P1 microbiomes (Figure 2c). As we similarly observed through ordination plots in Figure 1, the communities become more dissimilar to P1 over time. We then fit both a linear and quadratic regression to these data, and we found a better fit of a quadratic model than linear as evidenced by higher R^2^ and lower AIC values (Linear R^2^ 0.774, AIC -3563.231; Quadratic R^2^ 0.8379, AIC: -4414.637). Both models were highly significant (p<0.001). Taken together, this suggests that the rate of community change is slowing down, although it appears to have not entirely stopped.

We next observed changes in relative abundance of specific taxa within lines over time (Figure 2d, top 100 OTUs plotted). At each passage, there are numerous taxa that are differentially abundant compared to other passages. In some cases, there was evidence for replacement of OTUs within taxonomic groups. Specifically, in the top 10 most differentially abundant taxa as determined by using a Kruskal-Wallis test [27] (Supplemental Figure S3), three of them are in the family *Pseudomonadaceae*. Two *Pseudomonas* OTUs (0010, 0004) are in significantly higher relative abundance in P1 than in P4 (p<0.0001), and gradually decreased in relative abundance an unclassified *Pseudomonadaceae* (0002) is significantly more abundant in P4 as compared to other passages (p<0.0001). All three OTUs are present in the initial spray inoculum, although OTU0002 represents only 0.03% of rarified spray inoculum reads whereas *Pseudomonas* OTU0004 represents 27% and *Pseudomonas* OTU0010 represents 21%.

To better understand how bacterial community dynamics were changing over the course of the four passages, we utilized a recently developed cohesion metric to quantify connectivity of microbial community (Herren and McMahon 2017). In brief, community cohesion is a computational method used to predict within-microbiome dynamics by quantifying connectivity of microbial communities based on pairwise correlations and relative abundance of taxa. Changes in community cohesion over time are suggestive of biotic interactions, where connectivity can arise from either, or both, positive and negative interactions resulting from cross-feeding (positive) or competition (negative) as well as environmental co-filtering. When applied to our dataset (Supplemental Figure S4), we find a minor but significant increase in positive cohesion values (among 200 permutations) from P1 to P4 (R^2^=0.19, p=1.4 × 10^-38^). Consistent with positive cohesion values showing increased biotic interactions, there are also increasingly negative cohesion values from P1 to P4, which again is minor but significant (R^2^=0.257, p=1 × 10^-53^). To test our hypothesis that community change was due to deterministic and non-neutral processes, we first applied the Sloan neutral community model [28] and found that a neutral model is less correlated with observed communities on the plants over time (Supplemental Figure S5a). However, this model assumes equal dispersal amongst hosts, which was not the case for P2-P4, as microbiomes were passaged without pooling. Thus, we compared this finding to an approach that is more appropriate for our experimental design. We generated a null prediction based on the known community composition of inocula applied at each passage and comparing our observed communities to the predicted neutral community using a recently developed approached [29] (see methods for complete details). We found that Bray Curtis distances between predicted (null) and observed communities moderately increases over time (R^2^=0.261, p<0.0001) (Supplemental Figure S5b), suggesting that community change over the course of the passaging experiment is likely the result of deterministic rather than stochastic processes. Further evidence for a shift away from neutrality can be observed using occupancy-abundance curves in which the occupancy, or proportion of individuals in which an OTU is found, is plotted against its relative abundance. A positive correlation between the two is expected to occur by chance, as in a neutrally assembled community, but a change in distribution of individuals may indicate a community shaped by deterministic processes [30, 31]. When our data are visualized in this manner (Supplemental Figure S6), we see that in P1, the most abundant taxa also occupy the highest proportion of plants, as you would expect in a neutral community not undergoing niche selection. However, this trend collapses by P4 with many abundant taxa occupying far fewer individuals than would be expected under an assumption of neutrality.

We next designed an experiment to which we could apply Sloan’s model of neutral theory (Supplemental S7a). All lines from the end of P4 were pooled together and re-inoculated onto tomato plants, mimicking the inoculation procedure from the first passage. Plants that received the P4-combined inoculum had significantly different bacterial community composition than the P4 plants themselves (48% of variation explained, P=0.001; Supplemental S7b). We did not observe an effect of genotype on the communities assembled from this combined inoculum (p=0.565). We also found that the majority of the variation between samples (76%, p=0.001) was driven by an exceptional situation of introduction of a greenhouse taxon (OTU0003) to the plants (Supplemental S7c). To test if neutral processes were driving community structure in this experiment, we again examined fit to a neutral model using the Sloan model approach. In this case, as with P1, the assumption of equal dispersal potential among plants is met. In 200 iterative predictions, the fit of the neutral model is significantly higher in P1 (R^2^=0.87 ± 0.01) than P4 Combined (R^2^=0.52 ± 0.05; Student’s *t*-test, *p*-value < 0.01), suggesting that neutral processes are dictating the community structure after the first passage, but not in the P4 Combined experiment (Supplemental S7d). When P1 and P4 Combined are compared directly, we see the occupancy-abundance relationship breakdown in P4 Combined (Supplemental S7e).

### Mycobiome

For P1 and P4, we also used ITS amplicon sequencing to describe the fungal communities across lines, and observe patterns that are similar to the bacterial communities. We again found a significant effect of passage number on fungal communities (Figure 3a; Bray-Curtis distances for all samples, ADONIS, 43%, p=0.001). The significant effect of passage number remained after inoculum, control samples, and accession 2934 were removed (Figure 3b; 47%, p=0.001). However, unlike in the bacterial community analysis, we found no significant differences in community composition between control and experimental plants at P1 (p=0.117), P4 (p=0.649) or in both passages combined (p=0.588). Additionally, we found no effect of host genotype at either passage (p=0.612, p=0.576) or overall (p=0.997). We also measured a significant decrease in both OTU richness (t-test, p=0.013) and Shannon’s diversity (p=0.0005) between P1 and P4 across all genotypes (Figure 3c). Finally, analysis of the 5 most common taxa overall identified a single OTU, identified as *Rhodosporidiobolus nylandii*, which was not detectable in the inoculum or P1 but dominated the fungal community in P4 (Figure 3d).

**Figure 3.**
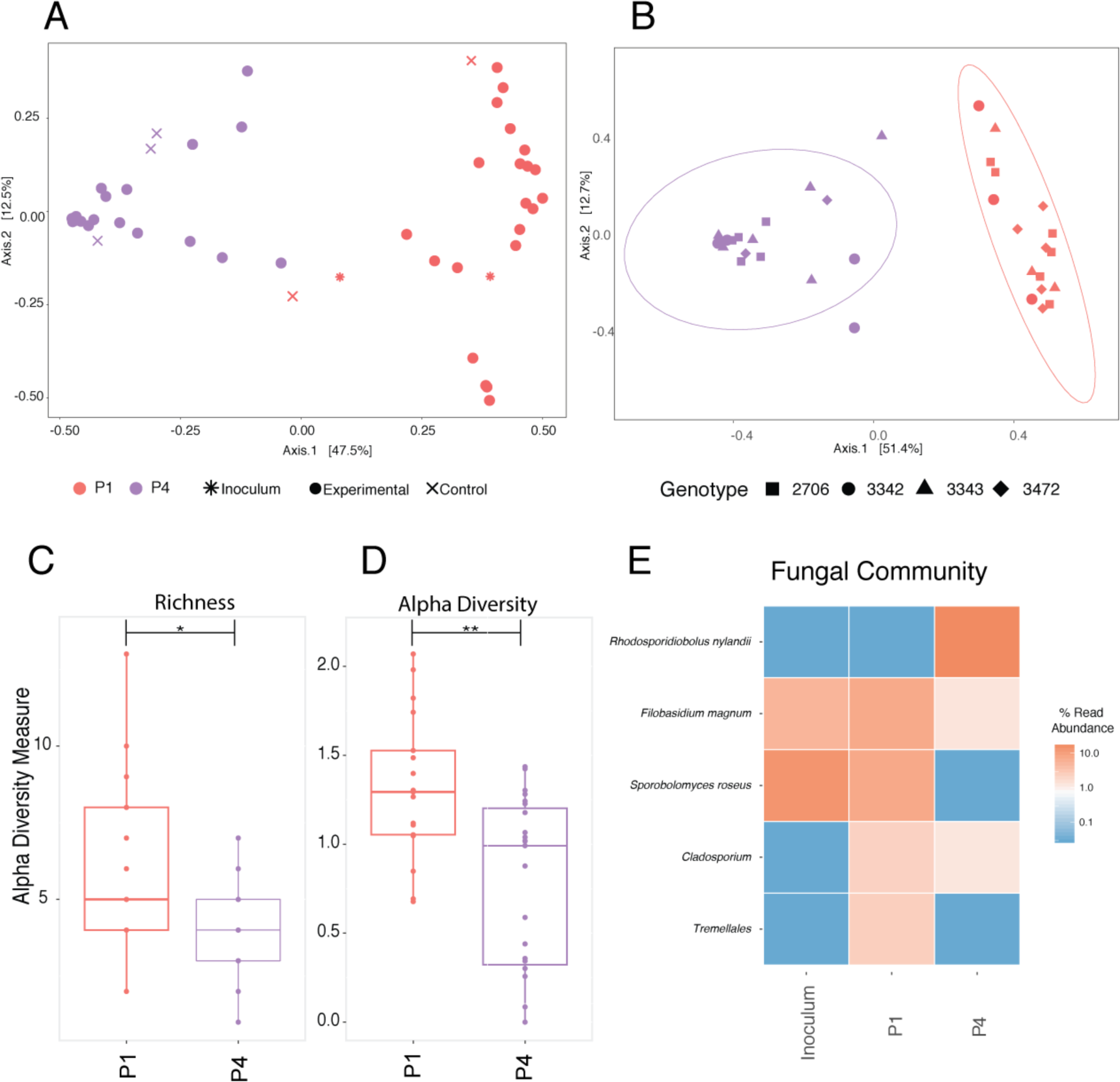
The Mycobiome. A PCoA plot of Bray-Curtis distances show a significant change in the community from P1 to P4, as determined by a PERMANOVA test (a). There is no effect of genotype (shapes) on the fungal community (b) Ellipses indicate 95% confidence around the clustering. Both richness (c) and Shannon’s alpha diversity (d) significantly decrease between P1 and P4. Relative abundance of the top five fungal taxa is plotted for the original inoculum, P1 and P4 (e). Significance values of pairwise comparisons in (c) and (d) are illustrated on the graph * p≤0.05; ** p≤0.01; *** p≤0.001; ****p≤0.0001.

### Testing microbiome adaptation using community coalescence

The similarity of changes in community structure both across replicates and genotypes over the course of the passaging experiment (Figures 1, 2, and 3) led us to predict that these microbiomes were becoming well adapted to the local plant conditions (by which we mean that the taxa present were positively selected for over time). To further determine if the community changes we observed from P1 to P4 were due to habitat selection rather than neutral processes, we employed a community coalescence competition experiment. In this experiment (Figure 4a), phyllosphere communities from the end of P1 (pooled across all lines) and the end of P4 (again, pooled across lines) were inoculated onto a new cohort of plants, either on their own or in an approximately 50:50 mixture of live cells (as determined using live/dead PMA treatment followed by ddPCR; see methods for complete details).

**Figure 4.**
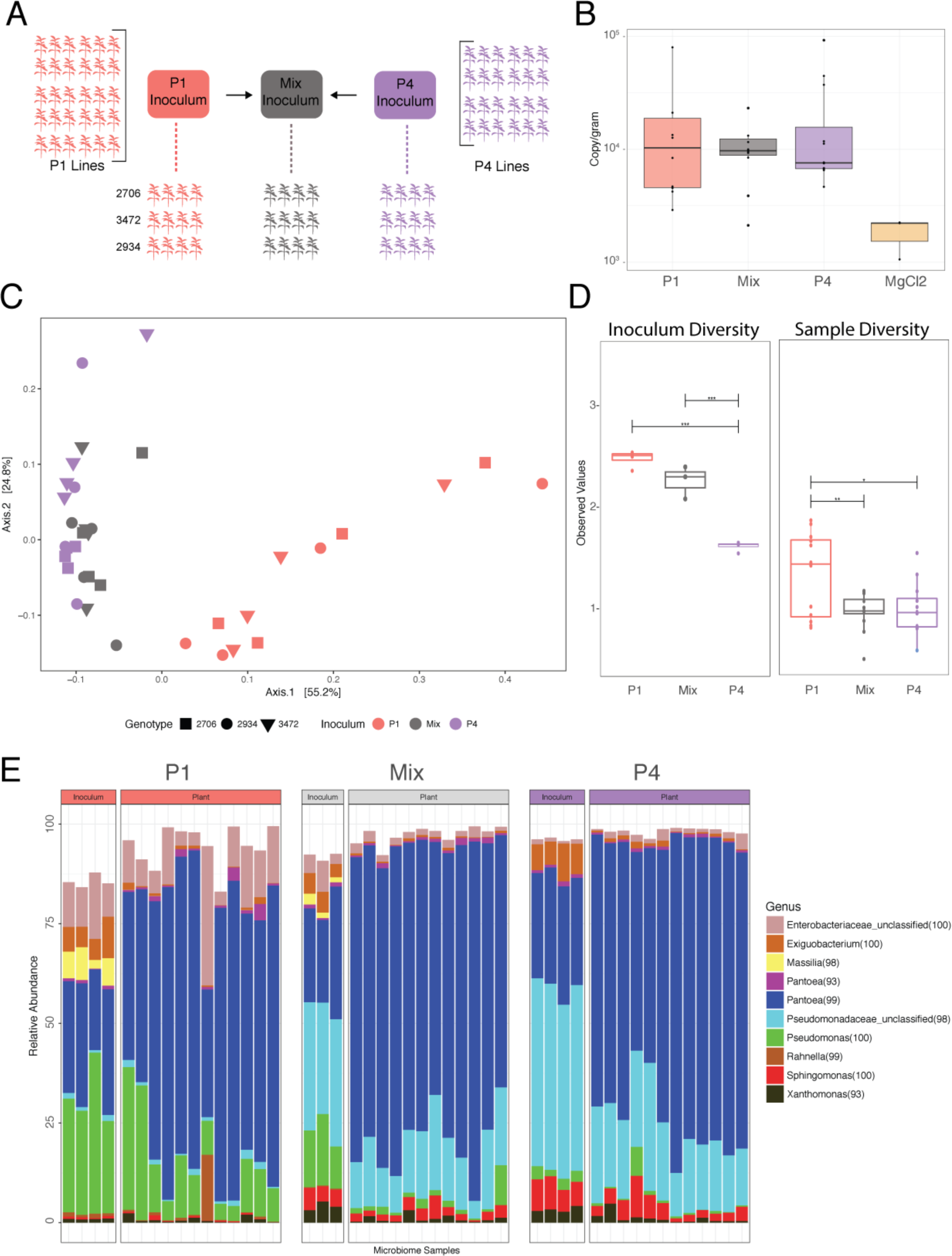
Testing microbiome adaptation. Plants were inoculated with pooled, passaged microbiomes from the end of P1, P4, or a 50:50 Mix of the two (a). Bacterial abundance was measured using ddPCR (b). A PCoA plot of Bray-Curtis distances colored by inoculum source shows that P1 plants have bacterial communities that are significantly different from P4 and Mix plants, which are indistinguishable (c). Shannon’s alpha diversity of the inoculum and experimental plants (d) show significant differences between samples. A bar graph illustrating composition of the top 10 OTUs shows differences in taxa amongst both the inoculum and experimental plants (e).

To ensure that our method for the mixed inoculum was effective, we sequenced multiple replicates of the P1, P4, and Mix inocula and found that source of inoculum explains 88% of dissimilarity amongst samples (ADONIS, p=0.002). To confirm that the Mix inoculum was significantly different than both P1 and P4 separately, we compared P1 and Mix inocula directly and found that 75% of difference between samples can be explained by this variable (p=0.02). Similarly, when P4 and Mix are compared directly, 74% of variation in the community is explained (p=0.02). This consistent difference among the three inocula allowed us to compare the communities colonizing plants from each treatment.

We first measured final bacterial abundance and found that colonization was lower on these plants than in previous experiments, but does not significantly differ among treatments (p=0.419), apart from control plants, where bacterial colonization was greatly reduced (Figure 4b). We then compared bacterial communities again using 16S amplicon sequencing and ordinated samples on a PCoA based on Bray-Curtis distances. Plants that received P1 inoculum have distinctly different communities than those that received either P4 or the Mixed inoculum. Plants that received the Mixed inoculum clustered together with those receiving P4 and were relatively indistinguishable. Using ADONIS tests, we determined that inoculum source can explain 45% of Bray-Curtis dissimilarity amongst samples (Figure 4c; p=0.001), and there was no effect of plant genotype (p=0.743; although note that only three genotypes were used in this experiment). In a pairwise analysis between P1 and Mixed, inoculum source explains 31% of the community dissimilarity (p=0.001). In contrast, inoculum source does not explain any significant variation in dissimilarity amongst P4 and Mixed inoculum plants (p=0.103). Together, these results suggest that the plants receiving the 50:50 mixed inoculum were indistinguishable in community composition from those receiving the pooled, P4 passaged microbiomes, and that these selected communities were not invadable by the microbial communities from the start of the experiment. Consistent with our results from the passaging experiment itself, alpha diversity was found to be highest in P1 plants compared to both P4 and Mixed plants (Figure 4d). Alpha diversity did not differ amongst communities colonizing plants from the P4 and Mixed inoculums, despite being different between the two inocula themselves. We also examined compositional makeup of the communities (Figure 4e), and consistent with P1 to P4 passaging results, we see differentially abundant taxa between groups (Supplemental Figure 8). Again, two *Pseudomonas* OTUs are more abundant in P1 plants as compared to P4 and Mix, in which there is an unclassified *Pseudomonaceae* that is higher in relative abundance.

## Discussion

The impact of a microbiome on host health and fitness depends not only on which microbial organisms are present in the community, but also on how they interact with one another within the microbiome [32]. Unlocking the great potential of microbiome manipulation and pre/probiotic treatment in reshaping host health will therefore depend on our ability to understand and predict these interactions. We took a microbiome passaging approach, inspired by classic experimental evolution, to test how selection for growth in the tomato phyllosphere under greenhouse conditions would impact microbiome diversity and adaptation across genotypes that differ in disease resistance genes.

Across independently selected lines passaged on five tomato genotypes, we observed a dramatic shift in community structure and composition, accompanied by a loss of alpha diversity (Figures 1 and 2). We also found that host genotype shapes community composition early in passaging (P1 and P2), explaining over 24% of variation amongst samples, but diminishes over time. The relative importance of host genotype and environment in shaping microbiome composition remains highly debated. Our results suggest that the relative importance of genotype versus other factors, such as the growth environment or strength of within-microbiome interactions, changes over the course of passaging on a constant host background. We observed that even in the absence of a strong genotype effect, there remains a legacy of genotype effect, in that OTUs found to be significantly associated with particular genotypes early on are more likely to be present at the end of passaging than those that did not exhibit any host preference.

In order to test if the phyllosphere microbiome undergoes habitat filtering, we chose to begin the experiment with a diverse inoculum. This starting community generated from field grown tomato plants likely contained microbes from other surrounding plant species, dust, soil, and other sources. In particular, neighboring plants have been shown to contribute to both the density and composition of local airborne microbes [33]. We found that although the total number of these field inoculum OTUs decreased over the course of the experiment, the taxa that remained consistently made up 78-95% of the community. This provides evidence that the original spray inoculum underwent strong niche selection over the course of the experiment. We also see evidence for niche selection through changing occupancy-abundance distributions. Gonzalez et al. found a similar breakdown of occupancy-abundance relations in animal communities using miniature moss microcosms [31]. The authors predict that this was due to dispersal limitation, as their experimental design created habitat fragmentation, and they did not observe this similar decline in correlation in communities that were connected by “habitat corridors”. In our experimental design, dispersal limitation is likely to have played a role in the changing community structure. In addition, the incidence of high-abundance, low-occupancy taxa in P4, or “clumping” [30], is further suggestive of niche selection.

To test the alternative hypothesis that community changes were due to neutral processes such as bottle necking or random dispersal, we first fit our data to neutral and null models, finding a poorer fit over time. We then tested this experimentally by conducting a community coalescence experiment to measure fitness of passaged microbiomes as compared to those from the start of the experiment. The results of this experiment strongly support the idea that these phyllosphere microbiomes adapted to the plant host environment over the course of four passages (Figure 4). Independent of overall bacterial abundance, P4 microbiomes were able to dramatically outcompete the less-adapted P1 microbiomes. This community coalescence approach [25] allowed us to demonstrate non-neutral selection of a bacterial community that is independent of host genotype and resistant to invasion by a more diverse, non-selected community. We cannot differentiate the relative contribution of evolutionary versus ecological change to the communities, but we expect both to have occurred within the time scale of these experiments. This community coalescence approach was used by others in a study conducted on methanogenic bacterial communities [34]. The authors found that when multiple methanogenic communities were combined, a single dominant community emerged from the mix. This emergent dominant community resembled the single community with the highest methane production that went into the combination, suggesting that the most-fit community is capable of reassembly, even in the presence of other community members.

While adaptation to both the local host environment (tomato plants, host genotype) and the larger environment (the greenhouse) were likely driving the increasingly non-neutral selection over time, the strength of within microbiome biotic interactions likely also increased over the course of the experiment. We see evidence for this through both increasing positive and negative community cohesion values. We also uncovered a strong effect of a greenhouse-acquired taxon on the community in one of the experiments (Figure S7). Though we are not able to determine what drove certain plants to be more colonized by this taxon than others, we did observe strong shifts in community composition associated with its relative abundance that may be due to spatial organization of plants in the greenhouse and/or stochastic initial colonization events. In a greenhouse study conducted on *Arabidopsis thaliana* phyllosphere communities, the authors found that abundance of certain dominant taxa could be tied to spatial organization of the plants that was likely driven by early stochastic events [13].

Although we focus primarily on the bacterial portion of the microbiome, the mycobiome changed over the course of passaging as well (Figure 3). Previous work in *A. thaliana* demonstrated that “hub” fungal taxa strongly influence both bacterial alpha and beta diversity [35]. Although it is possible that multi-kingdom interactions played a role in shaping community composition, our experimental methods, especially the process of sonicating epiphytic microbiota and freezing in between passages, likely biased passaging towards bacterial taxa and epiphytes. Similarly, pelleting of the community and removal of the supernatant at each passage would have selected against any free lytic bacteriophages. Previously, we found that the phage fraction of the microbiome is capable of altering both abundance and composition in the tomato phyllosphere [36]. Furthermore, our collection and inoculation method may have reduced selection for dispersal ability across the phyllosphere environment. By evenly spraying microbes onto leaves in a high humidity environment, we may have tipped the balance in favor of bacterial species that are better competitors within the microbiome. A dispersal-competition tradeoff was recently demonstrated using functional traits of soil microbial communities along a marine-to-land gradient, where bacterial communities from more disturbed habitats were found to be dominated by cell chemosensory and motility behaviors whereas those from more stable environments were dominated by traits for competition and chemical defense [37]. Future work is required to disentangle both the selective impacts of the plant versus environment versus multi-kingdom interactions in shaping microbiome adaptation, and the change in microbial function as a result of this response to selection.

Given the naturally distinct spatial structure, ease of sampling, high culturability, and demonstrated role in plant health [22, 38], the phyllosphere microbiome is an ideal model for testing theories of niche selection and microbiome adaptation. Through spray inoculation, the environment can be evenly saturated with diverse inoculum, and it is possible to sample the successfully colonized community its entirety. Moreover, bacterial abundance and growth can be tracked using ddPCR, and communities can be described using next generation sequencing. We were able to use the phyllosphere model to not only select upon entire host-associated microbial communities, but to then experimentally test our hypotheses regarding microbiome adaption in subsequent experiments. Using our approach, we also shed light on a notable challenge in microbiome research. Our data suggest that when describing the microbiome of an open environment, such as plant surfaces, many of the taxa found there may be transient visitors. In the case of the phyllosphere, there are microbes on leaf surfaces that may have emigrated from air, soil, surrounding plants, or other non-plant habitats and do not necessarily represent an adapted community that is capable of growth and persistence. Passaging of microbiomes in the absence of specific trait-based selection, as we have done here, seems to be a powerful way of differentiating those taxa that are, or can rapidly become, well adapted to the plant host environment. It also raises the question as to if a microbiome should be defined as the community that is found upon sampling and sequencing, or if a true microbiome is one that is adapted to its host or environment.

Overall, we were able to show rapid and robust habitat selection of these communities over relatively short time scales. The results uncover great promise of this approach and system for answering fundamental questions about the forces shaping microbiome assembly over time, and also pave the way for selecting stable, uninvadable host-associated microbiomes, which may inform rational microbiome manipulation and probiotic design. Experiments such as these are crucial if we are to understand general principles governing microbiome assembly and adaptation and use this knowledge for transformative applications in both medicine and agriculture.

## Materials/Methods (See supplement for complete methods)

### Tomato accessions

Tomato accessions were obtained from the Tomato Genetics Resource Center. Five tomato genotypes were used: *Solanum lycopersicum* money maker disease susceptible (TGRC 2706); *S. lycopersicum* money maker disease resistant (TGRC 3472); *S. lycopersicum* Rio Grande disease susceptible control for TGRC 3342 (TGRC 3343); *S. lycopersicum* Rio Grande disease resistant (TGRC 3342); and *S. pimpinellifolium* wild ancestor (2934). All genotypes were used for passages one, two, three, and p4-combined. Genotype 2934 was not used in passage four, as that genotype succumbed to fungal disease in the third generation. The community coalescence competition experiment included genotypes 2706, 3472, and 2934.

### Tomato germination and growth

Seeds were surface sterilized using TGRC recommendations then transferred onto 1% water agar plates and placed in the dark at 21°C until emergence of the hypocotyl. At that point, seedling plates were moved into a growth chamber and allowed to continue germination for 1 week. After approximately one week, seedlings were transferred planted in sunshine mix #1 soil in seedling trays. After approximately one more week of growth, seedlings were transplanted into 8” diameter pots, making the plants approximately 2.5-3 weeks old at the first time of microbial inoculation. Age of inoculation varied slightly from experiment to experiment but was kept identical amongst genotypes within an experiment.

### Inoculation preparation, first passage

Microbial inoculum for the first passage of the experiment was generated from field-grown tomato plants from the UC Davis Student Organic Farm collected in September and October of 2016. Above-ground plant material was collected from various genotypes of tomatoes across nine different sites spread through four fields. Other plant types, such as lettuce, eggplant, corn, and oak trees, surrounded the tomato fields. Sterile phosphate freezing buffer was added to the bags of leaves, and the entire bags were placed in a Branson M5800 sonicating water bath. Material was sonicated for 10 minutes. This gentle sonication washes microbes from the surfaces of the leaves but does not damage cells. The resulting leaf wash from each site was pooled and divided into 6 aliquots and stored in glycerol freezing buffer. For each inoculation in the first passage, an aliquot was thawed, cells pelleted, and re-suspended in 200mL 10mM MgCl_2_ buffer. Of this, 40mL were and heat killed in an autoclave for a 30 minutes at 121°C. Both live and heat-killed inoculum were plated. There was no growth from heat-killed inoculum, and live-inoculum concentration was calculated to be 1.1 ×’; 10 ^6 CFU/mL. Soil from each site, which had been stored at -20°C, was combined in a sterile bucket and thoroughly mixed before inoculation.

### Inoculation procedure

Soil inoculation: The top layer of every pot was supplemented with 40 grams of UC Davis Farm Soil. Soil inoculation was only performed once and only for the first passage of plants. Spray inoculation: Each plant was sprayed with 4.5mL of inocula using misting spray tops. Control plants from passage 1 were inoculated with the heat-killed inocula. Control plants from P2 onward were inoculated with sterile 10mM MgCl_2_. Immediately after inoculation, plants were placed in a random order in a high-humidity misting chamber for 24 hours. After 24 hours, the plants were moved to a greenhouse bench. Plants were inoculated once per week in the same manner and were placed in the misting chamber for 24 hours after every inoculation.

### Plant sampling and inoculation preparation for P2-4 (Figures 1, 2, and 3)

Ten days after the final spray inoculation, plants were sampled. With the exception for plant cohort 5, all plants were cut off at the base and immediately placed into sterile 1L bottles individually. By the end of cohort 5, the plants had grown too large to sample the entire plant, and instead, roughly 2/3 of the plant material was sampled from each plant, with care taken to sample the same age of branches from every plant. After collection, plant material was weighed, sterile buffer added, and the entire bottle sonicated as above. Half of the volume from each plant was pelleted and re-suspended in ∼1mL of 1:1 KB Broth Glycerol and stored at -80°C for inoculation of the subsequent passage. The other half of the volume was pelleted and stored as a pellet at -20°C for DNA extractions. To prepare inoculation of the next passage, microbiome glycerol stocks were thawed, briefly pelleted to remove glycerol, and re-suspended in sterile 10mM MgCl_2_.

### Inoculation preparation, combination of P4 microbiomes (Figure S7)

Frozen microbiomes from all plants from the end of passage four were thawed, and half the volume was removed from each aliquot. These aliquots were combined into one pooled meta-inoculum. This was divided into six aliquots. One was used immediately, and the rest of the aliquots were stored at -20°C in KB Glycerol and thawed by aliquot for each week of inoculation, as above.

### P1, P4 coalescence experiment (Figure 4)

Genotypes 2706, 3472, and 2934 were used for this experiment, and four plants of each genotype received each treatment (P1, P4, and Mix). One control plant of each genotype was spray inoculated with MgCl_2_ as a control. To prepare the inoculum, microbiomes from the end of passage one and the end of passage four were combined. The same was done for all of the individual microbiomes that came off of passage 4 plants. In order to quantify only live cells, we used PMA treatment, using a method adapted from others [39], prior to ddPCR quantification (see below). Bacterial concentration was matched to 7.7 × 10^6 cells/mL. Plants were inoculated for three weeks and harvested 10 days after the final inoculation as described previously.

### Bacterial quantification using ddPCR

The BioRad QX200 system was used for culture independent quantification of bacteria. Complete ddPCR methods are described elsewhere [36]. Bacterial abundance was measured directly after microbes were sonicated off plant surfaces into sterile buffer. For consistency, the same region of the 16S gene used below for amplicon sequencing was used for bacterial quantification. PNAs were used as well to limit any background amplification of plant mitochondrial or chloroplast DNA. All data were normalized to weight, in grams, and concentrations are reported as 16S copy number/gram.

### DNA extractions

DNA was extracted from microbial pellets using the Qiagen PowerSoil DNA extraction kit. A buffer control extraction was included for every set of extractions in order to identify and exclude taxa present in the dataset due to buffer contamination.

### 16S Libraries

The 16S rRNA gene was amplified using dual-indexed primers designed for the V3-V4 region [40] using the following primers: 341F (5 -CCTACGGGNBGCASCAG-3) and 785R (5 -GACTACNVGGGTATCTAATCC-3) [41]. Additionally, we also used peptide nucleic acids, PNAs [42] to decrease amplification of plant mitochondrial and chloroplast DNA. Negative buffer controls and PCR controls were sequenced along with experimental samples. Amplicons from each sample were pooled in equimolar concentrations, cleaned using an AMpure bead clean-up kit. Libraries were prepared for paired 300-nucleotide reads in Illumina’s MiSeq V3 platform (Illumina) at The California Institute for Quantitative Biosciences (QB3) at UC Berkeley and run in 1 lane.

### ITS Libraries

Using the same DNA as above, the ITS2 region was amplified using ITS9-F: GAACGCAGCRAAIIGYGA and ITS4-R: TCCTCCGCTTATTGATATGC following a protocol published online by the Joint Genome Institute. A second PCR was performed (7 cycles) in order to anneal MiSeq illumunia adapters and barcodes onto the amplicons. PCRs were carried out in duplicate and pooled before they were prepared for sequencing by the QB3 sequencing facility as described above.

### Data Processing and Analysis

MiSeq sequencing files were demultiplexed by QB3 sequencing facility. Bacterial reads were combined into contigs using VSearch [43], and the remainder of the analysis was carried out in Mothur [44] following their MiSeq SOP [45] (See supplement for specifics). We used a 97% similarity cut-off for defining OTUs and the Silva reference database [46] for taxonomic assignment. Bacterial were rarified to 8,000 reads per sample. For the fungal community, an OTU table was generated from the fungal community sequencing data using QIIME 2 (version 2018.8) (See supplement for specifics). Reads were clustered into OTUs at 97% identity and assigned taxonomy using the UNITE database and the feature-classifier plug-in [47]. Once bacterial and fungal OTU tables were generated in Mothur and QIIME, the remainder of the analysis was performed in R using the following packages: Phyloseq [48], vegan [49], ampvis2 [50], and MicrobiomeSeq (Alfred Ssekagiri, William T. Sloan, Umer Zeeshan Ijaz).

### Community Cohesion Metrics

The estimations of positive and negative cohesion values follows the cohesion metrics approach proposed by Herren *et al*. [51]. We modified their method to estimate cohesion values by using two relative abundance profiles of a training set and test set. Relative abundance profile of the training set was obtained by randomly selecting half of the samples in each microbiome passage. The test set consists of the other half of the samples. Using the training set and following the same procedure as Herren *et al*., connectedness metrics were calculated. The estimated connectedness metrics subtracts a null model. The obtained connectedness metrics are multiplied by relative abundance profile of test set to estimate positive and negative cohesion values. Two hundred iterations of sampling randomization in each microbiome passage were carried out at OTU level to obtain training set and test set for P1, P2, P3, and P4.

### Neutral model

The neutral model was proposed by Sloan *et al*. to describe both microbial diversity and taxa-abundance distribution of a community [28]. Burns *et al*. [16] have developed a R package based on Sloan’s neutral model to determine the potential importance of neutral process to a community assembly. In brief, the neutral model creates a potential neutral community by a single free parameter describing the migration rate, *m*, based on two sets of abundance profiles – a local community and metacommunities. The local community describes the observed relative abundance of OTUs, while the metacommunity is estimated by the mean relative abundance across all local communities. The estimated migration rate is the probability of OTU dispersal from the metacommunity to replace a randomly lost individual in the local community. The migration rate can be interpreted as dispersal limitation. In each microbiome passage, half of the samples were randomly selected and the relative abundance profile at the OTU level was used. The neutral model fit and migration rate were estimated in the resolution results of 200 iterations for P1, P2, P3, P4, and P4 Combined.

### Null model predictions

We applied a null model approach on the serial passaging data P1-P4 to characterize the changes of stochastic process driving the assembly of plant microbiome over time. Lines that had high quality sequencing data at every time point (thirteen in total) were used for this analysis. The null scenario for each line at each passage was generated using the data for that same line at the previous passage. The null scenario of P1 was generated using the original field inoculum sample. The null model approach was based on community pairwise dissimilarity proposed by Chase and Myers [52] and extended by Stegen *et al*. to incorporate species abundance [53]. Chase and Myers proposed a degree of species turnover by a randomization procedure where species probabilistically occur at each local community until observed local richness is reached. However, the estimated degree of turnover does not include species abundance. To take full advantage of our dataset, we also incorporated species relative abundance into the procedure proposed by Stegen *et al*. Zinger *et al*. has developed R code for the null model and applied the null model approach on the soil microbiome [29]. This approach does not require *a priori* knowledge of the local community condition and determines if each plant microbiome at the current passage deviates from a null scenario generated by that same microbiome at the previous passage. In brief, the null scenario of each was generated by random resampling of OTUs and remained the same richness and number of reads with the original sample. Total OTUs observed in the sample and the corresponding relative abundance were used as probabilities of selecting an OTU and its associated number of reads, respectively. The Bray-Curtis distance is used to calculate dissimilarities across null communities with 1,000 permutations. The average of dissimilarities among permutations represents null expectations of community dissimilarities. The null deviation shows the differences between average null expectation and the observed microbiome of the same line.

## Acknowledgements

The authors would like to acknowledge the UC Davis Student Farm, who provided access to the fields from which the original inoculum was generated. They would also like to thank Christina Winstrom and the Oxford Tract greenhouse staff for their role in plant care throughout the experiments. The authors thank Shirley Zhang for assistance with plant inoculation. Lastly, the authors thank Dylan Smith and Shana McDevitt for their continued support with sequencing efforts for the experiment.

