## Supplemental for "Successive passaging of a plant-associated microbiome reveals robust habitat and host genotype-dependent selection"

Title

* To whom correspondences should be addressed

Table 1……………………………………………………………...Page 10

Complete Methods……………………………………………....Page 13

References……………………………………………………......Page 24

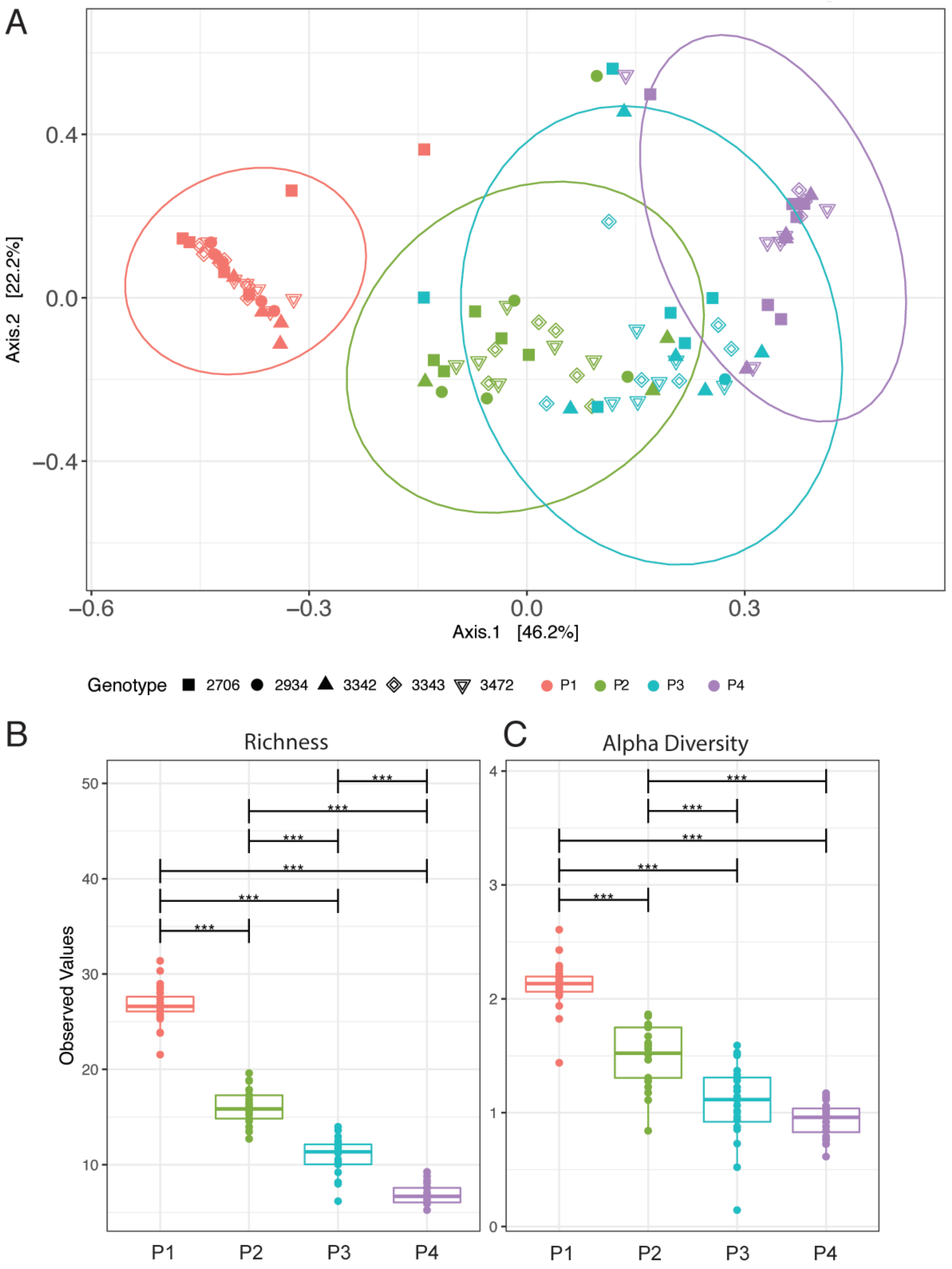

**Figure S1 Serial passaging of the phyllosphere microbiome: Inoculum only taxa**
The dataset was subsampled to only contain OTUs that were present in the initial spray inoculum. A PCoA plot of Bray-Curtis distances of experimental plants shows a significant effect (determined by PERMANOVAs) of passage (colors) and genotype (shapes) (a). Richness (b) and Shannon's alpha diversity (c) of each experimental plant are plotted at each passage and show a significant decrease over time. Significance values of pairwise comparisons are illustrated on the graph * $p \leq 0.05$; ** $p \leq 0.01$; *** $p \leq 0.001$; **** $p \leq 0.0001$.

A

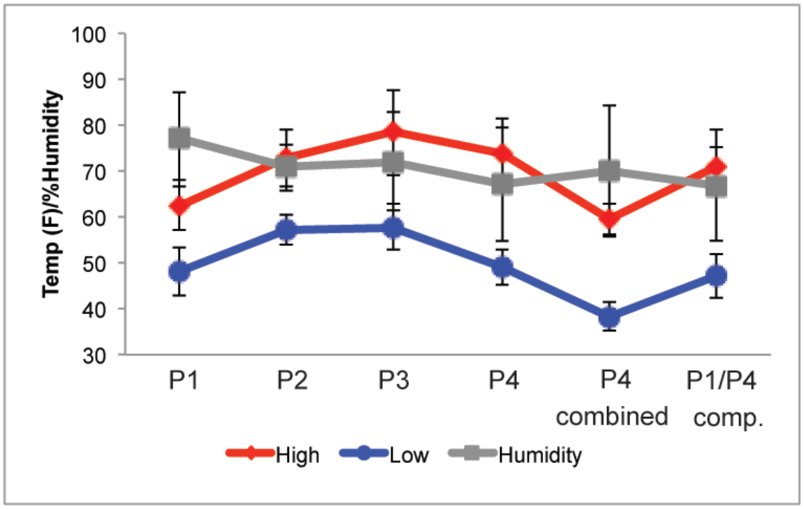

B

| Passage/Exp | Start | End |
|---|---|---|
| One | 11/8/16 | 12/15/16 |
| Two | 6/1/17 | 7/4/17 |
| Three | 8/25/17 | 9/26/17 |
| Four | 10/3/17 | 11/1/17 |
| P4 Combined | 11/28/17 | 1/3/18 |
| P1/P4 Comp. | 10/18/18 | 11/12/18 |

**Figure S2 Climatic Variables**
Although all experiments were performed in the greenhouse, outside climatic variables varied for the duration of the six experiments. Humidity, high temp, and low temp are plotted (a). The date at which the plants were first inoculated and the date at which they were harvested are shown (b).

A

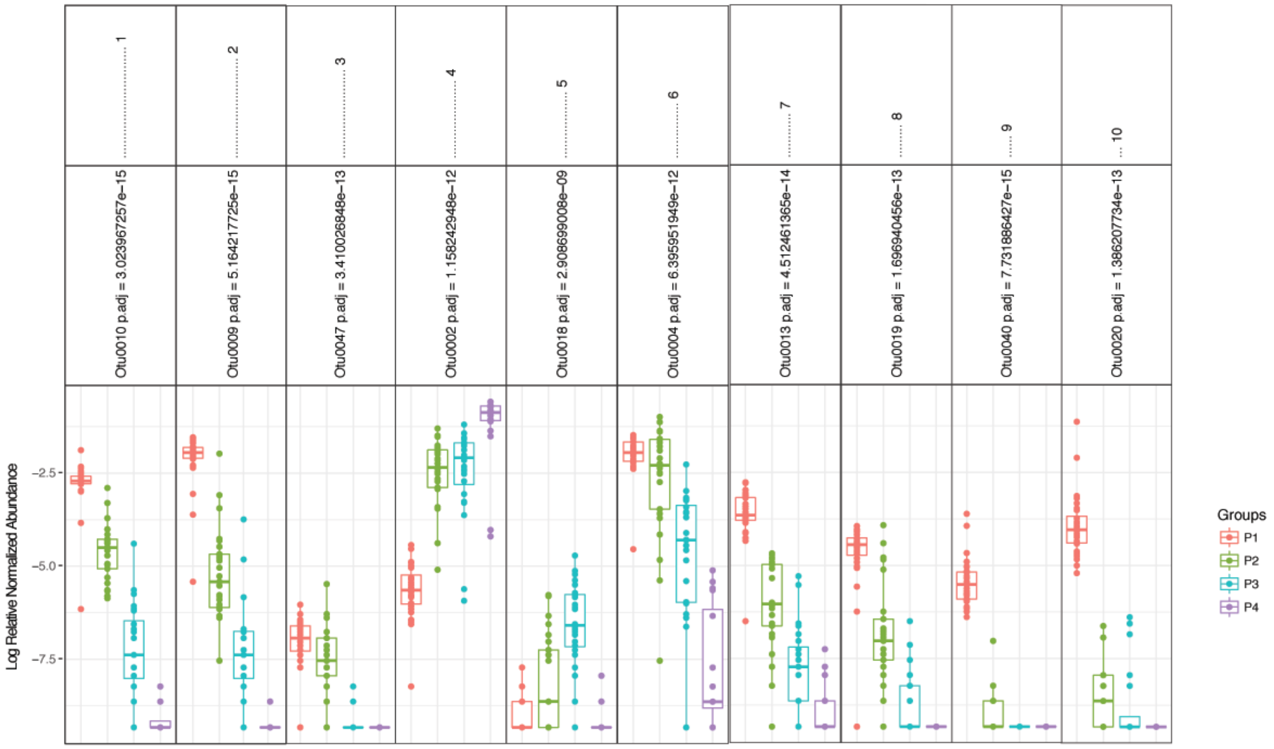

B

| | Phylum | Class | Order | Family | Genus |
|---|---|---|---|---|---|
| **Otu0010** | Proteobacteria | Gammaproteobacteria | Pseudomonadales | Pseudomonadaceae | Pseudomonas |
| **Otu0009** | Proteobacteria | Gammaproteobacteria | Enterobacteriales | Enterobacteriaceae | Enterobacteriaceae_unclassified |
| **Otu0047** | Firmicutes | Bacilli | Bacillales | Family_XII | Exiguobacterium |
| **Otu0002** | Proteobacteria | Gammaproteobacteria | Pseudomonadales | Pseudomonadaceae | Pseudomonadaceae_unclassified |
| **Otu0018** | Proteobacteria | Betaproteobacteria | Burkholderiales | Oxalobacteraceae | Massilia |
| **Otu0004** | Proteobacteria | Gammaproteobacteria | Pseudomonadales | Pseudomonadaceae | Pseudomonas |
| **Otu0013** | Proteobacteria | Betaproteobacteria | Burkholderiales | Oxalobacteraceae | Massilia |
| **Otu0019** | Proteobacteria | Gammaproteobacteria | Enterobacteriales | Enterobacteriaceae | Rahnella |
| **Otu0040** | Bacteroidetes | Sphingobacteriia | Sphingobacteriales | Sphingobacteriaceae | Pedobacter |
| **Otu0020** | Bacteroidetes | Sphingobacteriia | Sphingobacteriales | Sphingobacteriaceae | Pedobacter |

**Figure S3 Differentially abundant taxa amongst passaged lines**
We performed a Kruskal-Wallis test on log-relative transformed OTU abundance at different passages using the MicrobiomeSeq package (a). This is a non-parametric method, and it tests whether samples originate from the same distribution. P-values are corrected for multiple testing using family wise error rates. Significant OTU rankings 1-10 are assigned importance using random forest classifier. Identities of OTUs are displayed as well (b).

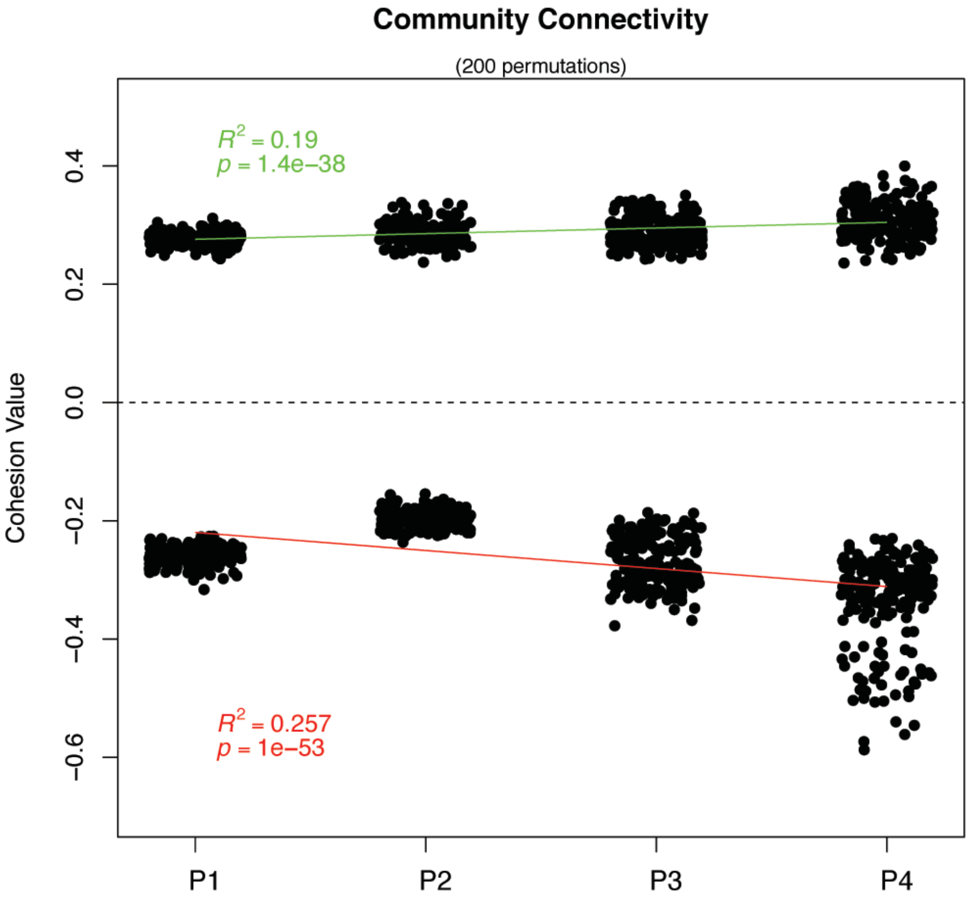

**Figure S4 Community Cohesion from P1 to P4**
We applied community cohesion metrics on our dataset to describe microbial dynamics in P1, P2, P3, and P4. We calculated both positive and negative cohesion values and found a mild but significant increase in positive and negative cohesion values from P1 to P4.

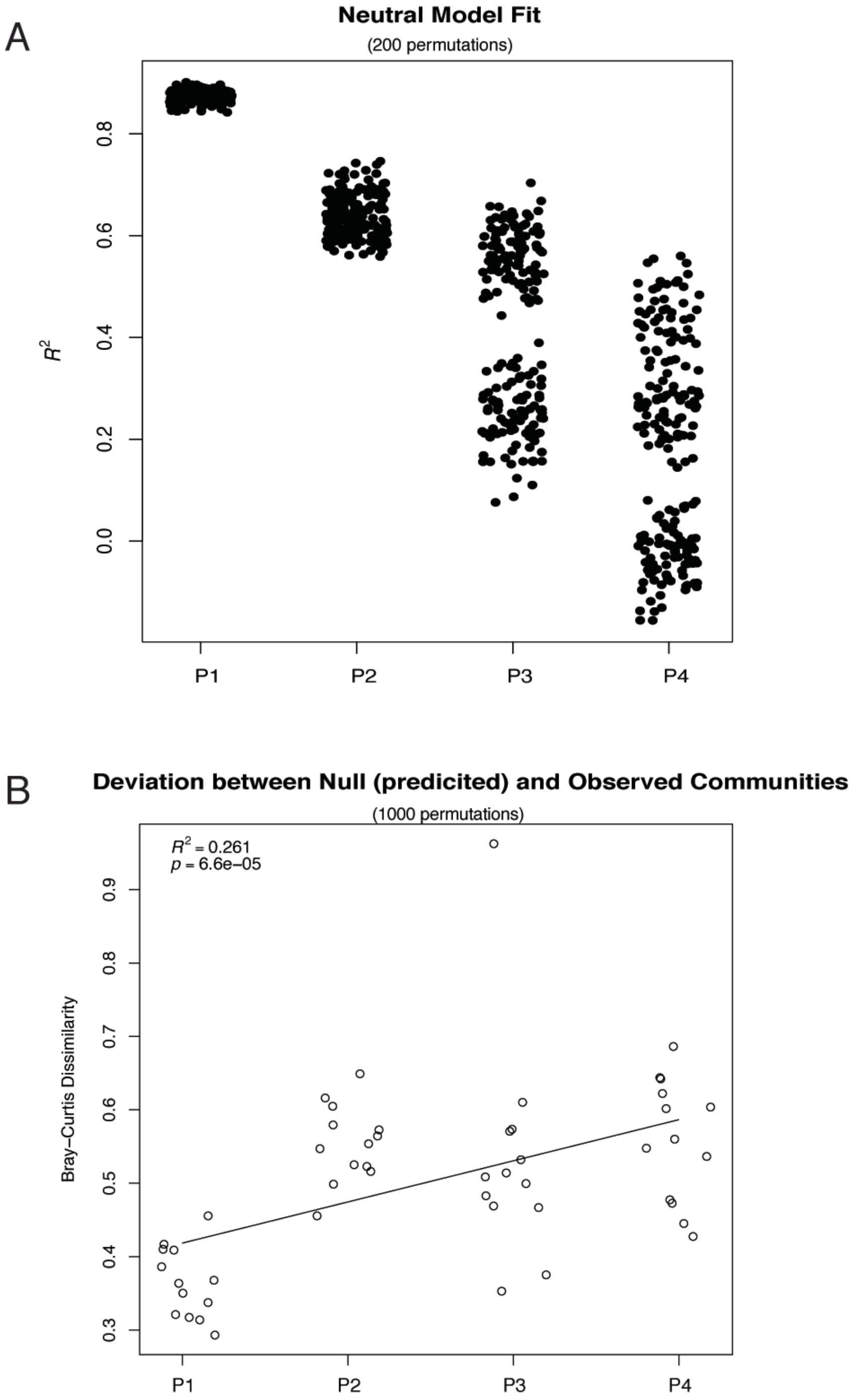

**Figure S5 Neutral model fit and comparison of predicted null to observed communities** We compared the observations of community structure and predicted community composition using a neutral model and found a poorer fit over time (a). We compared Bray-Curtis distances between a predicted null model from the n-1 passage with observed communities from passage n over 1,000 iterations and we found a mild but significant increase in deviation from the null prediction from P1 to P4 (b).

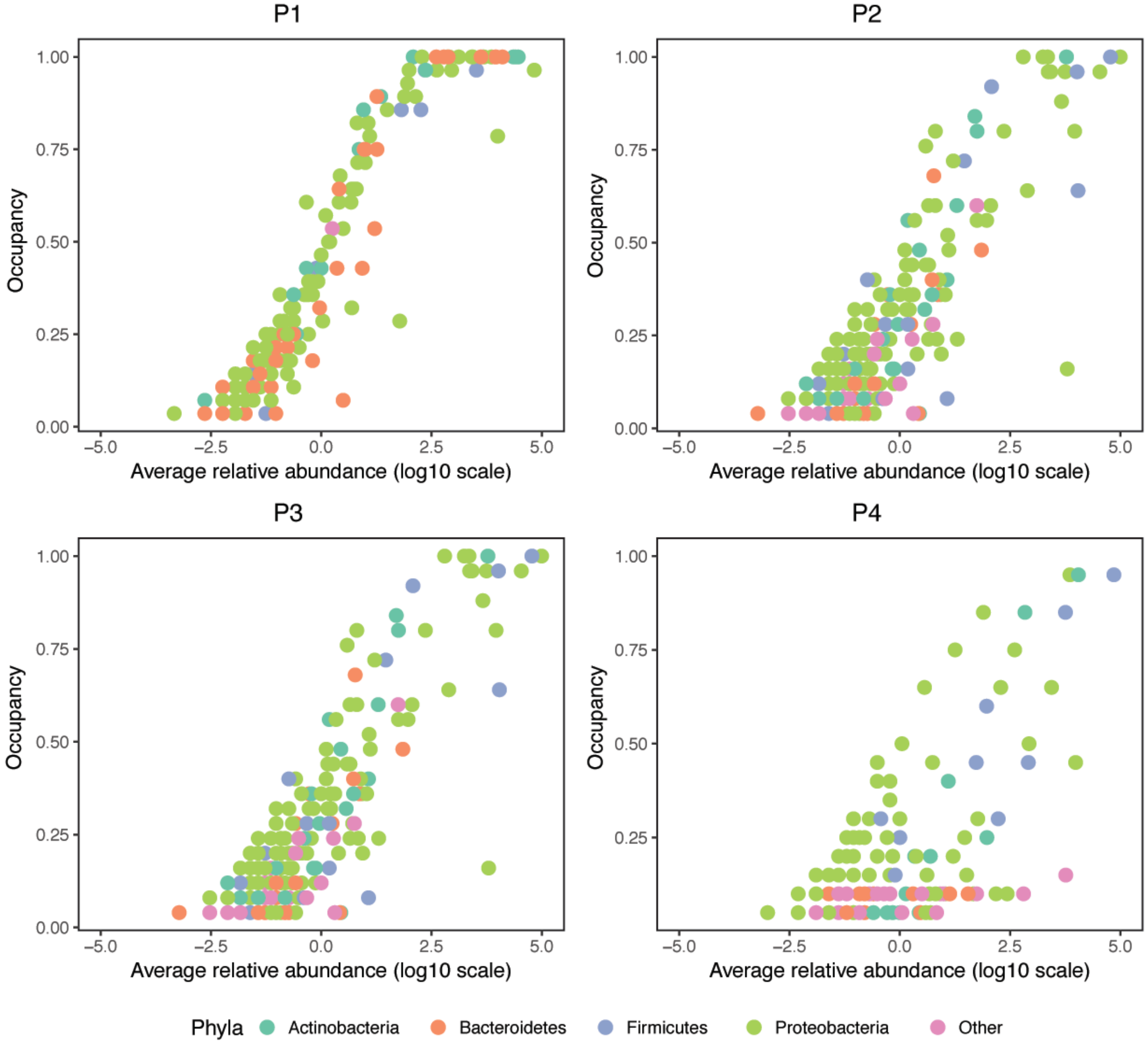

**Figure S6 Occupancy- Abundance relationships of taxa from P1 to P4**
For each OTU, its occupancy (or, proportion of plant hosts in which it was found) is plotted against the log 10 of its relative abundance. OTUs belonging phyla other than those in the top 4 Phyla are classified as “other”.

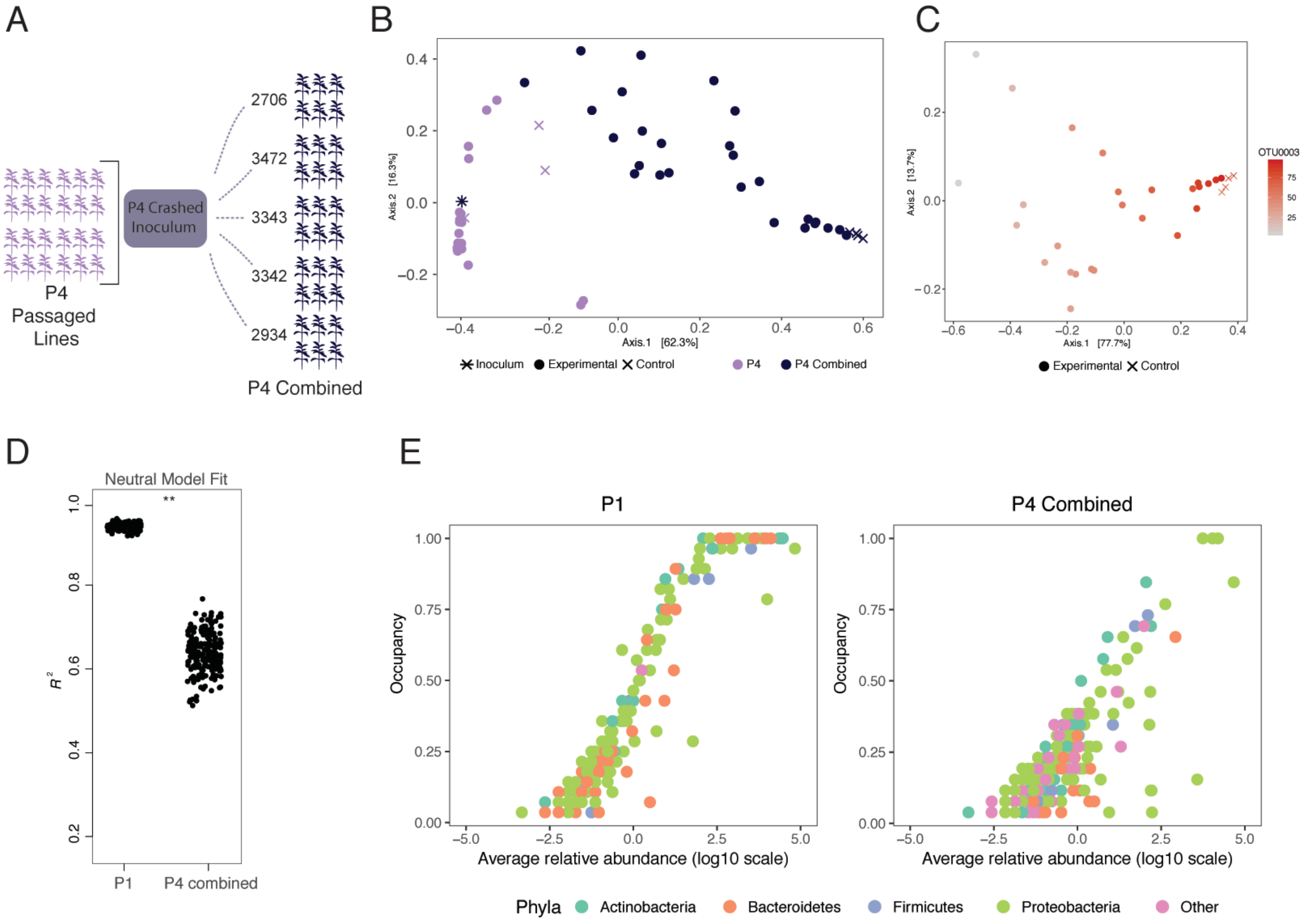

**Figure S7 Combination of passaged lines and re-incoulation**
Passaged microbiomes from the end of P4 were pooled and then sprayed onto a fifth cohort of tomato plants (six replicates of five genotypes) (a). P4 plants, inoculum generated from P4 plants, and controls are plotted on a PCoA plot based on Bray-Curtis distances (b), and P4-Combined plants cluster apart from P4 plants. We compared the observations of community structure and predicted community composition by a neutral model for P1 and P4-Combined (c), and in 200 iteratively predictions, the fit of the neutral model is significantly higher in P1 than P4-Combined (Student's *t*-test, *p*-value < 0.01). Visualized on a PCoA plot (d), relative abundance of OTU0003 can be seen driving Bray-Curtis distances amongst plants, both experimental and control. Occupancy-Abundance curves for P1 and P4-combined are shown side-by-side (e).

A

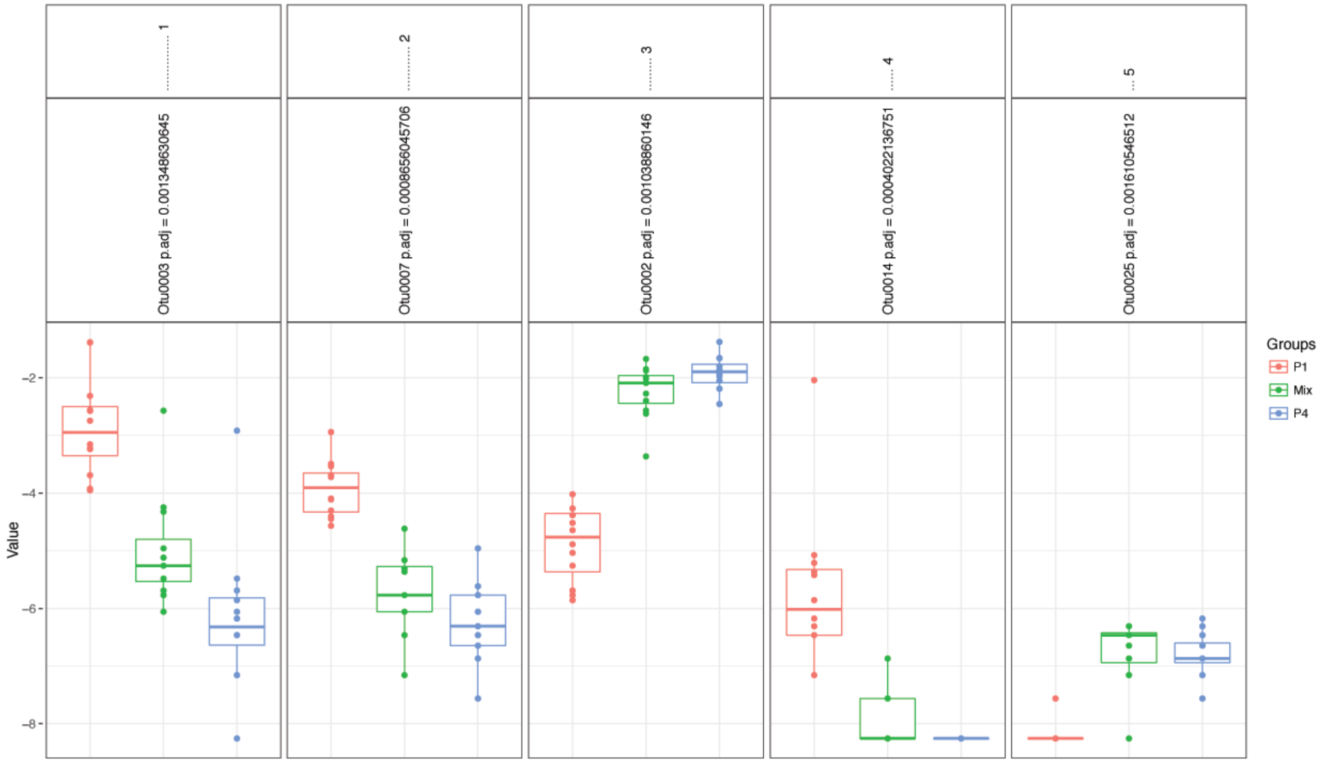

B

| | Phylum | Class | Order | Family | Genus |
|---|---|---|---|---|---|
| Otu0003 | Proteobacteria | Gammaproteobacteria | Pseudomonadales | Pseudomonadaceae | Pseudomonas |
| Otu0007 | Proteobacteria | Gammaproteobacteria | Pseudomonadales | Pseudomonadaceae | Pseudomonas |
| Otu0002 | Proteobacteria | Gammaproteobacteria | Pseudomonadales | Pseudomonadaceae | Pseudomonadaceae_unclassified |
| Otu0014 | Proteobacteria | Gammaproteobacteria | Enterobacteriales | Enterobacteriaceae | Rahnella |
| Otu0025 | Proteobacteria | Gammaproteobacteria | Enterobacteriales | Enterobacteriaceae | Enterobacteriaceae_unclassified |

**Figure S8 Differentially abundant taxa in community coalescence experiment**
We performed a Kruskal-Wallis test on log-relative transformed OTU abundance at different passages using the MicrobiomeSeq package (a). This is a non-parametric method, and it tests whether samples originate from the same distribution. P-values are corrected for multiple testing using family wise error rates. Significant OTU rankings 1-5 are assigned importance using random forest classifier. Identities of OTUs are displayed as well (b).

### Supplemental Table 1: Top 100 OTU taxonomic assignments

| OTU | Phylum | Class | Order | Family | Genus |
|---|---|---|---|---|---|
| 0001 | Proteobacteria(100) | Gammaproteobacteria(100) | Enterobacteriales(100) | Enterobacteriaceae(100) | Pantoea(98) |
| 0002 | Proteobacteria(100) | Gammaproteobacteria(100) | Pseudomonadales(100) | Pseudomonadaceae(100) | Unclassified(93) |
| 0003 | Proteobacteria(100) | Betaproteobacteria(100) | Burkholderiales(100) | Oxalobacteraceae(100) | Unclassified(97) |
| 0004 | Proteobacteria(100) | Gammaproteobacteria(100) | Pseudomonadales(100) | Pseudomonadaceae(100) | Pseudomonas(100) |
| 0005 | Proteobacteria(100) | Alphaproteobacteria(100) | Sphingomonadales(100) | Sphingomonadaceae(99) | Sphingomonas(99) |
| 0006 | Proteobacteria(100) | Gammaproteobacteria(100) | Xanthomonadales(100) | Xanthomonadaceae(100) | Xanthomonas(100) |
| 0008 | Proteobacteria(100) | Gammaproteobacteria(100) | Pseudomonadales(100) | Pseudomonadaceae(100) | Pseudomonas(100) |
| 0009 | Proteobacteria(100) | Gammaproteobacteria(100) | Enterobacteriales(100) | Enterobacteriaceae(100) | Unclassified(100) |
| 0010 | Proteobacteria(100) | Gammaproteobacteria(100) | Pseudomonadales(100) | Pseudomonadaceae(100) | Pseudomonas(100) |
| 0011 | Proteobacteria(100) | Gammaproteobacteria(100) | Enterobacteriales(100) | Enterobacteriaceae(100) | Unclassified(98) |
| 0012 | Firmicutes(100) | Bacilli(100) | Bacillales(100) | Family_XII(100) | Exiguobacterium(100) |
| 0013 | Proteobacteria(100) | Betaproteobacteria(100) | Burkholderiales(100) | Oxalobacteraceae(100) | Massilia(99) |
| 0014 | Proteobacteria(100) | Gammaproteobacteria(100) | Enterobacteriales(100) | Enterobacteriaceae(100) | Unclassified(100) |
| 0015 | Proteobacteria(100) | Alphaproteobacteria(100) | Caulobacterales(100) | Caulobacteraceae(100) | Brevundimonas(100) |
| 0016 | Firmicutes(100) | Bacilli(100) | Bacillales(100) | Bacillaceae(100) | Bacillus(100) |
| 0017 | Proteobacteria(100) | Alphaproteobacteria(100) | Sphingomonadales(100) | Sphingomonadaceae(100) | Novosphingobium(100) |
| 0018 | Proteobacteria(100) | Betaproteobacteria(100) | Burkholderiales(100) | Oxalobacteraceae(100) | Massilia(100) |
| 0019 | Proteobacteria(100) | Gammaproteobacteria(100) | Enterobacteriales(100) | Enterobacteriaceae(100) | Rahnella(56) |
| 0020 | Bacteroidetes(100) | Sphingobacteriia(100) | Sphingobacteriales(100) | Sphingobacteriaceae(100) | Pedobacter(100) |
| 0021 | Proteobacteria(100) | Gammaproteobacteria(100) | Pseudomonadales(100) | Pseudomonadaceae(100) | Pseudomonas(100) |
| 0022 | Proteobacteria(100) | Gammaproteobacteria(100) | Xanthomonadales(100) | Xanthomonadaceae(100) | Stenotrophomonas(100) |
| 0023 | Proteobacteria(100) | Gammaproteobacteria(100) | Xanthomonadales(100) | Xanthomonadaceae(100) | Stenotrophomonas(99) |
| 0024 | Actinobacteria(100) | Actinobacteria(100) | Micrococcales(100) | Microbacteriaceae(100) | Curtobacterium(100) |
| 0025 | Firmicutes(100) | Bacilli(100) | Bacillales(100) | Bacillaceae(100) | Bacillus(100) |
| 0026 | Actinobacteria(100) | Actinobacteria(100) | Micrococcales(100) | Microbacteriaceae(100) | Unclassified(100) |
| 0027 | Proteobacteria(100) | Alphaproteobacteria(100) | Rhizobiales(100) | Methylobacteriaceae(100) | Methylobacterium(100) |
| 0028 | Proteobacteria(100) | Gammaproteobacteria(100) | Enterobacteriales(100) | Enterobacteriaceae(100) | Unclassified(67) |
| 0029 | Proteobacteria(100) | Betaproteobacteria(100) | Burkholderiales(100) | Oxalobacteraceae(100) | Massilia(100) |
| 0032 | Proteobacteria(100) | Alphaproteobacteria(100) | Rhizobiales(100) | Methylobacteriaceae(100) | Methylobacterium(100) |
| 0033 | Actinobacteria(100) | Actinobacteria(100) | Micrococcales(100) | Sanguibacteraceae(100) | Sanguibacter(100) |
| 0034 | Actinobacteria(100) | Actinobacteria(100) | Micrococcales(100) | Microbacteriaceae(100) | Unclassified(91) |
| 0035 | Proteobacteria(100) | Alphaproteobacteria(100) | Rhizobiales(100) | Methylobacteriaceae(100) | Methylobacterium(100) |
| 0036 | Proteobacteria(100) | Betaproteobacteria(100) | Burkholderiales(100) | Burkholderiaceae(100) | Ralstonia(100) |
| 0037 | Proteobacteria(100) | Gammaproteobacteria(100) | Enterobacteriales(100) | Enterobacteriaceae(100) | Pantoea(97) |
| 0038 | Proteobacteria(100) | Gammaproteobacteria(100) | Enterobacteriales(100) | Enterobacteriaceae(100) | Unclassified(98) |
| 0039 | Bacteroidetes(100) | Sphingobacteriia(100) | Sphingobacteriales(100) | Chitinophagaceae(100) | Sediminibacterium(100) |
| 0040 | Bacteroidetes(100) | Sphingobacteriia(100) | Sphingobacteriales(100) | Sphingobacteriaceae(100) | Pedobacter(100) |
| 0041 | Bacteroidetes(100) | Sphingobacteriia(100) | Sphingobacteriales(100) | Sphingobacteriaceae(100) | Pedobacter(100) |
| 0042 | Proteobacteria(100) | Betaproteobacteria(100) | Burkholderiales(100) | Oxalobacteraceae(100) | Duganella(100) |

| | | | | | |
|---|---|---|---|---|---|
| 0043 | Firmicutes(100) | Bacilli(100) | Bacillales(100) | Bacillaceae(100) | Bacillus(99) |
| 0045 | Firmicutes(100) | Bacilli(100) | Bacillales(100) | Paenibacillaceae(100) | Paenibacillus(100) |
| 0046 | Actinobacteria(100) | Actinobacteria(100) | Streptomycetales(100) | Streptomycetaceae(100) | Streptomyces(100) |
| 0047 | Firmicutes(100) | Bacilli(100) | Bacillales(100) | Family_XII(100) | Exiguobacterium(100) |
| 0048 | Proteobacteria(100) | Alphaproteobacteria(100) | Rhizobiales(100) | Rhizobiaceae(100) | Rhizobium(59) |
| 0049 | Chloroflexi(100) | Ktedonobacteria(100) | Ktedonobacterales(100) | Unclassified(79) | Unclassified(79) |
| 0050 | Actinobacteria(100) | Actinobacteria(100) | Streptomycetales(100) | Streptomycetaceae(100) | Streptomyces(76) |
| 0052 | Proteobacteria(100) | Alphaproteobacteria(100) | Sphingomonadales(100) | Unclassified(98) | Unclassified(98) |
| 0053 | Proteobacteria(100) | Gammaproteobacteria(100) | Enterobacteriales(100) | Enterobacteriaceae(100) | Unclassified(90) |
| 0055 | Proteobacteria(100) | Alphaproteobacteria(100) | Rhizobiales(100) | Methylobacteriaceae(100) | Methylobacterium(100) |
| 0056 | Proteobacteria(100) | Gammaproteobacteria(100) | Pseudomonadales(100) | Pseudomonadaceae(100) | Pseudomonas(100) |
| 0057 | Bacteroidetes(100) | Flavobacteriia(100) | Flavobacteriales(100) | Flavobacteriaceae(100) | Chryseobacterium(100) |
| 0059 | Proteobacteria(100) | Gammaproteobacteria(100) | Enterobacteriales(100) | Enterobacteriaceae(100) | Unclassified(100) |
| 0061 | Proteobacteria(100) | Gammaproteobacteria(100) | Pseudomonadales(100) | Pseudomonadaceae(100) | Unclassified(83) |
| 0062 | Bacteroidetes(100) | Flavobacteriia(100) | Flavobacteriales(100) | Flavobacteriaceae(100) | Chryseobacterium(100) |
| 0065 | Proteobacteria(100) | Alphaproteobacteria(100) | Sphingomonadales(100) | Sphingomonadaceae(100) | Unclassified(94) |
| 0066 | Firmicutes(100) | Bacilli(100) | Bacillales(100) | Paenibacillaceae(100) | Paenibacillus(100) |
| 0068 | Actinobacteria(100) | Actinobacteria(100) | Micrococcales(100) | Micrococcaceae(100) | Pseudarthrobacter(94) |
| 0069 | Proteobacteria(100) | Gammaproteobacteria(100) | Pseudomonadales(100) | Pseudomonadaceae(100) | Pseudomonas(100) |
| 0070 | Proteobacteria(100) | Gammaproteobacteria(100) | Enterobacteriales(100) | Enterobacteriaceae(100) | Unclassified(98) |
| 0071 | Firmicutes(100) | Bacilli(100) | Bacillales(100) | Paenibacillaceae(100) | Paenibacillus(100) |
| 0073 | Proteobacteria(100) | Alphaproteobacteria(100) | Rhizobiales(100) | Bradyrhizobiaceae(100) | Bradyrhizobium(68) |
| 0074 | Proteobacteria(100) | Alphaproteobacteria(100) | Caulobacterales(100) | Caulobacteraceae(100) | Caulobacter(100) |
| 0076 | Actinobacteria(100) | Actinobacteria(100) | Micrococcales(100) | Microbacteriaceae(100) | Rathayibacter(98) |
| 0077 | Proteobacteria(100) | Gammaproteobacteria(100) | Enterobacteriales(100) | Enterobacteriaceae(100) | Unclassified(98) |
| 0079 | Proteobacteria(100) | Gammaproteobacteria(100) | Pseudomonadales(100) | Moraxellaceae(100) | Acinetobacter(100) |
| 0081 | Bacteroidetes(100) | Sphingobacteriia(100) | Sphingobacteriales(100) | Sphingobacteriaceae(100) | Pedobacter(100) |
| 0082 | Proteobacteria(100) | Gammaproteobacteria(100) | Xanthomonadales(100) | Xanthomonadaceae(100) | Koukoulia(100) |
| 0083 | Bacteroidetes(100) | Sphingobacteriia(100) | Sphingobacteriales(100) | Sphingobacteriaceae(100) | Sphingobacterium(100) |
| 0085 | Proteobacteria(100) | Gammaproteobacteria(100) | Enterobacteriales(100) | Enterobacteriaceae(100) | Unclassified(75) |
| 0086 | Bacteroidetes(100) | Flavobacteriia(100) | Flavobacteriales(100) | Flavobacteriaceae(100) | Flavobacterium(100) |
| 0088 | Proteobacteria(100) | Alphaproteobacteria(100) | Rhodospirillales(100) | Acetobacteraceae(100) | Roseomonas(100) |
| 0089 | Proteobacteria(100) | Gammaproteobacteria(100) | Enterobacteriales(100) | Enterobacteriaceae(100) | Unclassified(96) |
| 0093 | Proteobacteria(100) | Gammaproteobacteria(100) | Pseudomonadales(100) | Pseudomonadaceae(100) | Pseudomonas(86) |
| 0094 | Proteobacteria(100) | Betaproteobacteria(100) | unclassified(99) | Unclassified(99) | unclassified(99) |
| 0096 | Bacteroidetes(100) | Sphingobacteriia(100) | Sphingobacteriales(100) | Sphingobacteriaceae(100) | Pedobacter(100) |
| 0097 | Firmicutes(100) | Bacilli(100) | Bacillales(100) | Paenibacillaceae(100) | Saccharibacillus(100) |
| 0098 | Proteobacteria(100) | Alphaproteobacteria(100) | Caulobacterales(100) | Caulobacteraceae(100) | Brevundimonas(100) |
| 0106 | Actinobacteria(100) | Actinobacteria(100) | Streptosporangiales(100) | Thermomonosporaceae(100) | Actinoallomurus(100) |
| 0115 | Actinobacteria(100) | Actinobacteria(100) | Micrococcales(100) | Microbacteriaceae(100) | Curtobacterium(92) |
| 0116 | Proteobacteria(100) | Betaproteobacteria(100) | Burkholderiales(100) | Alcaligenaceae(100) | Verticia(100) |

| | | | | | |
|---|---|---|---|---|---|
| 0118 | Proteobacteria(100) | Gammaproteobacteria(100) | Xanthomonadales(100) | Xanthomonadaceae(100) | Stenotrophomonas(100) |
| 0123 | Proteobacteria(100) | Gammaproteobacteria(100) | Enterobacteriales(100) | Enterobacteriaceae(100) | Unclassified(94) |
| 0126 | Proteobacteria(100) | Alphaproteobacteria(100) | Caulobacterales(100) | Caulobacteraceae(100) | Unclassified(82) |
| 0130 | Actinobacteria(100) | Actinobacteria(100) | Micrococcales(100) | Micrococcaceae(100) | Paenarthrobacter(67) |
| 0134 | Proteobacteria(100) | Alphaproteobacteria(100) | Rhodospirillales(100) | Incertae Sedis(100) | Reyranella(100) |
| 0136 | Proteobacteria(100) | Gammaproteobacteria(100) | Pseudomonadales(100) | Pseudomonadaceae(100) | Pseudomonas(80) |
| 0150 | Proteobacteria(100) | Alphaproteobacteria(100) | Rhizobiales(100) | Methylobacteriaceae(100) | Methylobacterium(100) |
| 0167 | Actinobacteria(100) | Actinobacteria(100) | Micrococcales(100) | Micrococcaceae(100) | Arthrobacter(95) |
| 0169 | Actinobacteria(100) | Actinobacteria(100) | Frankiales(100) | Sporichthyaceae(100) | Unclassified(100) |
| 0174 | Proteobacteria(100) | Betaproteobacteria(100) | Burkholderiales(100) | Comamonadaceae(100) | Acidovorax(96) |
| 0189 | Proteobacteria(100) | Gammaproteobacteria(65) | Pseudomonadales(65) | Pseudomonadaceae(65) | Unclassified(65) |
| 0190 | Armatimonadetes(100) | Fimbriimonadia(100) | Fimbriimonadales(100) | Fimbriimonadaceae(100) | Unclassified(99) |
| 0208 | Bacteroidetes(100) | Sphingobacteriia(100) | Sphingobacteriales(100) | Chitinophagaceae(100) | Heliimonas(100) |
| 0213 | Bacteroidetes(100) | Flavobacteriia(100) | Flavobacteriales(100) | Flavobacteriaceae(100) | Epilithonimonas(99) |
| 0215 | Bacteroidetes(100) | Sphingobacteriia(100) | Sphingobacteriales(100) | Sphingobacteriaceae(100) | Pedobacter(100) |
| 0222 | Bacteroidetes(100) | Sphingobacteriia(100) | Sphingobacteriales(100) | Chitinophagaceae(100) | Vibrionimonas(100) |
| 0223 | Bacteroidetes(100) | Sphingobacteriia(100) | Sphingobacteriales(100) | Sphingobacteriaceae(100) | Pedobacter(100) |
| 0293 | Proteobacteria(100) | Alphaproteobacteria(100) | Rhodospirillales(100) | Acetobacteraceae(100) | Unclassified(100) |
| 0323 | Bacteroidetes(100) | Sphingobacteriia(100) | Sphingobacteriales(100) | Chitinophagaceae(100) | Sediminibacterium(100) |

Complete Methods

**Tomato accessions**

Tomato accessions were obtained from the Tomato Genetics Resource Center. Five tomato genotypes were used: *Solanum lycopersicum* money maker disease susceptible (TGRC 2706); *S. lycopersicum* money maker disease resistant (TGRC 3472); *S. lycopersicum* Rio Grande disease susceptible control for TGRC 3342 (TGRC 3343); *S. lycopersicum* Rio Grande disease resistant (TGRC 3342); and *S. pimpinellifolium* wild ancestor (2934). During the 2016 growing season, seeds for these experiments were generated by growing tomatoes in sunshine mix soil in the Jane Gray Greenhouse at UC Berkeley. Sunshine Mix #1 soil was used. Upon flowering, plants were manually pollinated by flicking flowers. Care was taken to switch gloves between plants of different genotypes. Fruits were collected and placed in plastic Ziploc bags, manually crushed, and allowed to ferment at 21°C for 2-3 weeks. After the fermentation process was complete, seeds were strained from remaining fruit material, rinsed with DI water, and allowed to dry on filter paper. Seeds were stored in the dark at 21°C until use. All genotypes were used for passages one, two, three, and p4- combined. Genotype 2934 was not used in passage four, as that genotype succumbed to fungal disease in the third generation. The community coalescence competition experiment included genotypes 2706, 3472, and 2934.

**Tomato germination and growth**

Seeds were surface sterilized using TGRC recommendations as follows: seeds were soaked in 2.7% bleach (sodium hypochlorite) solution for 20 minutes. Sterilized seeds were then washed with sterile $ddH_2O$ three times to remove any excess bleach. Sterilized seeds were then transferred onto 1% water agar plates and placed in the dark at 21°C until emergence of the hypocotyl. At that point, seedling plates were moved into a growth chamber and allowed to continue germination for 1 week. Growth chamber conditions were 25°C, 65% humidity and 16 h daylight per day. After approximately one week, seedlings were transferred planted in sunshine mix #1 soil in seedling trays. After approximately one more week of growth, seedlings were transplanted into 8” diameter pots, making the plants approximately 2.5-3 weeks old at the first time of microbial inoculation. Age of inoculation varied slightly from experiment to experiment but was kept identical amongst genotypes within an experiment.

**Inoculation preparation, first passage**

Microbial inoculum for the first passage of the experiment was generated from field-grown tomato plants from the UC Davis Student Organic Farm collected in September and October of 2016. One-gallon Ziploc bags were filled with leaf, stem, and some flower material from tomato plants. One bag was collected from each of nine different sites, spread through four different fields. Plant material was collected from various genotypes of tomatoes. Other plant types, such as lettuce, eggplant, corn, and oak trees, surrounded the tomato fields. During the October collection, soil was also collected at each site. The top ~2cm of soil was brushed away, and a 50mL conical was pushed directly into the soil at the base of a plant which was in the middle of each collection site. Plant material and soil were transferred to the lab on ice and stored at 4°C briefly until processing. Sterile phosphate freezing buffer was added to the bags of leaves, and the entire bags were placed in a Branson M5800 sonicating water bath. Material was sonicated for 10 minutes. This gentle sonication washes microbes from the surfaces of the leaves but does not damage cells.  The resulting leaf wash from each site was pooled. From the September collection, leaf wash was pelleted for 10 mins at 4000 x G, re-suspended in glycerol freezing buffer, and stored at -80 for approximately one month. This was then thawed, re-spun to remove the freezing buffer, and combined with the October leaf wash. At that point, the start inoculum was divided into 6 aliquots and stored in glycerol freezing buffer. For each inoculation in the first passage, an aliquot was thawed and cells pelleted for 10 mins at 4000 X G. Cells were re-suspended in 200mL 10mM $MgCl_2$ buffer. Of this, 40mL were and heat killed in an autoclave for a 30 minutes at 121°C. Inoculum was plated, and an absence of growth confirmed that the heat-kill was effective. To get initial concentration of inoculum, dilution plating was performed on Kings Broth agar plates (1.1 X 10 ^6 CFU/mL). Soil from each site, which had been stored at -20°C, was combined in a sterile Nalgene bucket and thoroughly mixed before inoculation.

**Inoculation procedure**

Soil inoculation: The top layer of every pot was supplemented with 40 grams of UC Davis Farm Soil. Soil inoculation was only performed once and only for the first passage of plants. Spray inoculation: Each plant was sprayed, using misting spray tops placed in 15mL conicals, with approximately 4.5mL of inocula. Control plants from passage 1 were inoculated with the heat-killed inocula. Control plants from subsequent experiments were inoculated with sterile 10mM $MgCl_2$. Immediately after inoculation, plants were placed in a random order in a high-humidity misting chamber for 24 hours. After 24 hours, the plants were moved to a greenhouse bench. Plants were inoculated once per week in the same manner and were placed in the misting chamber for 24 hours after every inoculation. Passage one plants received 5 weeks of inoculation, P2-P4: four weeks, and the fifth cohort: five weeks.

**Plant sampling and inoculation preparation for P2-4 (Figures 1, 2, and 3)**

Ten days after the final spray inoculation, plants were sampled. With the exception for plant cohort 5, all plants were cut off at the base and immediately placed into sterile 1L bottles individually. By the end of cohort 5, the plants had grown too large to sample the entire plant, and instead, roughly 2/3 of the plant material was sampled from each plant, with care taken to sample the same age of branches from every plant. After collection, plant material was weighed, and 200mL of sterile 10mM $MgCl_2$ were added to each bottle containing the plant material. The bottles were submerged in a sonicating water bath, sonicated for 5 minutes, vortexed, and sonicated for another 5 minutes. Half of the volume from each plant was pelleted for 10 mins at 4200 X G, re-suspended in ~1mL of 1:1 KB Broth Glycerol, divided into aliquots, and stored at -80°C for inoculation of the subsequent passage. The other half of the volume was pelleted in the same manner and then stored as a pellet at -20°C for DNA extractions. To prepare inoculation of the next passage, microbiome glycerol stocks were thawed, briefly pelleted to remove glycerol, and re-suspended in sterile 10mM $MgCl_2$. Volume of re-suspension depended slightly on the size of the plants, but in general ranged from 5-10mL. Microbiomes were never pooled.

**Inoculation preparation, combination of P4 microbiomes (Figure S7)**

Frozen microbiomes from all plants from the end of passage four were thawed, and half the volume was removed from each aliquot. These aliquots were combined into one pooled meta-inoculum. This was divided into six aliquots. One was used immediately, and the rest of the aliquots were stored at -20°C in KB Glycerol and thawed by aliquot for each week of inoculation, as above.

**P1, P4 coalescence experiment (Figure 4)**

Genotypes 2706, 3472, and 2934 were used for this experiment, and four plants of each genotype received each treatment (P1, P4, and Mix). One control plant of each genotype was spray inoculated with $MgCl_2$ as a control. To prepare the inoculum, microbiomes from the end of passage one and the end of passage four were used. All aliquots (one from each plant, except for plant 4 which had exhibited disease symptoms) were thawed and combined. The same was done for all of the individual microbiomes that came off of passage 4 plants. To remove the glycerol, the samples were spun down and re-resuspended in 10mM $MgCl_2$. In order to generate the 50/50 mix of P1 and P4 microbiomes, live/dead PCR with PMA treatment was used, adapted from the following method (Carini et al. 2017). Briefly, serial dilutions of P1 and P4 were performed in $MgCl_2$. Each sample then received PMA at a final concentration of 100uM and vortexed. Samples were incubated in the dark at room temp for 5 minutes. Then they were placed in ice on a tray exactly 10cm away from a 700 watt halogen lamp. The light was turned on for 30 seconds, and turned off for 30 seconds. During the 30 seconds without light, the samples were all vortexed. This was repeated three more times. Samples were then pelleted for 10 minutes at 5000 X G. The supernatant including the excess PMA was removed, and cells were re-suspended in sterile 10mM $MgCl_2$. Droplet Digital PCR (as described below) was then utilized to quantify bacteria from each sample, and concentration was matched to 7.7 x 10^6 cells/mL. P1 and P4 were aliquotted separately and then re-combined for the mixed inoculum so that each plant received ~9 x 10^4 bacteria each week that they were inoculated. Plants were inoculated for three weeks and harvested 10 days after the final inoculation as described previously.

**Bacterial quantification using ddPCR**

The BioRad QX200 system was used for culture independent quantification of bacteria. Complete ddPCR methods are described elsewhere (Morella et al. 2018). Bacterial abundance was measured directly after microbes were sonicated off plant surfaces into sterile buffer. For consistency, the same region of the 16S gene used below for amplicon sequencing was used for bacterial quantification.  PNAs were used as well to limit any background amplification of plant mitochondrial or chloroplast DNA. Five ul of sample were used in every reaction. All data were normalized to weight, in grams, and concentrations are reported as 16S copy number/gram.

**DNA extractions**

DNA was extracted from microbial pellets using the Qiagen PowerSoil DNA extraction kit. A buffer control extraction was included for every set of extractions in order to identify and exclude taxa present in the dataset due to buffer contamination.

**16S Libraries**

The 16S rRNA gene was amplified using dual-indexed primers designed for the V3- V4 region (Naylor et al. 2017) using the following primers: 341F (5 -CCTACGGGNBGCASCAG-3) and 785R (5 -GACTACNVGGGTATCTAATCC-3) (Takahashi et al. 2014). Additionally, we also used peptide nucleic acids, PNAs (Lundberg et al. 2013) to decrease amplification of plant mitochondrial and chloroplast DNA. Negative buffer controls and PCR controls were sequenced along with experimental samples. Reaction conditions were 94°C for 3 min, 94°C for 45 s, 78°C for 10 s, 50°C for 1 min, 72°C for 1.5 min, repeat steps 2–5 30 times and 72°C for 10 min. PCR mixtures were randomized in order, run in duplicate for each sample, pooled and quantified using Qubit. Amplicons from each sample were pooled in equimolar concentrations, cleaned using an AMpure bead clean-up kit. Libraries were prepared for paired 300-nucleotide reads in Illumina's MiSeq V3 platform (Illumina) at The California Institute for Quantitative Biosciences (QB3) at UC Berkeley and run in 1 lane.

**ITS Libraries**

Using the same DNA as above, the ITS2 region was amplified using ITS9-F: GAACGCAGCRAAIIGYGA and ITS4-R: TCCTCCGCTTATTGATATGC following a protocol published online by the Joint Genome Institute. A second PCR was performed (7 cycles) in order to anneal MiSeq illumunia adapters and barcodes onto the amplicons. PCRs were carried out in duplicate and pooled before they were prepared for sequencing by the QB3 sequencing facility as described above.

**Data Processing and Analysis**

MiSeq sequencing files were demultiplexed by QB3 sequencing facility. Reads were combined into contigs using VSearch (Rognes et al. 2016), and the remainder of the analysis was carried out in Mothur (Schloss et al. 2009) following their MiSeq SOP (Kozich et al. 2013). Data were quality-filtered, and chimeras were removed using UChime (Edgar et al. 2011). Singletons were removed using the split.abund command in Mothur after pre-clustering of similar sequences. We used a 97% similarity cut-off for defining OTUs. The Silva reference database (Quast et al.

2013) was used for sequence alignment and taxonomic assignment. Archaeal, chloroplast, mitochondrial and unknown domain DNA sequences were removed. To account for reagent contaminants, we also sequenced two DNA extraction kit controls and PCR controls along with our samples. Contaminant OTUs from control samples that were at a similar or higher relative abundance in control samples compared to experimental samples were removed from the full OTU table. Bacterial were rarified to 8,000 reads per sample. For the fungal community, an OTU table was generated from the fungal community sequencing data using QIIME 2. Trimmed, paired reads were first denoised, without read trimming, using the DADA2 plug-in (Callahan et al. 2016). Chimeric sequences were then filtered using the uchime-denovo command of the Vsearch plug-in (Rognes et al. 2016). Reads were then clustered into OTUs at 97% identity using the cluster-features-closed-reference command in the VSEARCH plug-in and the 2017 version of the UNITE database (Nilsson et al. 2019). In order to assign taxonomy to the clustered OTUs, a Naïve-Bayes classifier was first trained using the UNITE database and the feature-classifier plug-in (Bokulich et al. 2018). The classify-sklearn command of the feature-classifier plug-in was finally used to assign taxonomy to the clustered OTUs. Once bacterial and fungal OTU tables were generated in Mothur and QIIME, the remainder of the analysis was performed in R using the following packages: Phyloseq (McMurdie and Holmes 2013), vegan (Dixon and Palmer 2003), ampvis2 (Skytte Andersen et al. 2018), and MicrobiomeSeq (Alfred Ssekagiri, William T. Sloan, Umer Zeeshan Ijaz). For determining the effects of specific variables on Bray-Curtis dissimilarities between samples, PERMANOVA tests were run using Vegan's Adonis and adonis2 functions. For the genotype effect observed in P1 and P2, the data were also analyzed with the removal of the primary outlying line in P1, which was a diseased plant. The significance does not change (30% of dissimilarity explained, p=0.002). That same line had too low read depth to be analyzed at P2, and thus was excluded from this analysis at the rarefaction step. By P3, this line was included, as it did not fall outside of the 95% confidence intervals for P3 clustering. Goodness of fit for linear and quadratic models of Bray-Curtis distance change was based on both $R^2$ values and calculating Akaike information criterion values.

**Community Cohesion Metrics**

The estimations of positive and negative cohesion values follows the cohesion metrics approach proposed by Herren *et al*. (Herren and McMahon 2017). Herren *et al*. multiplied the connectedness metrics determined by relative abundance profile by the same relative abundance profile to estimate cohesion values. We modified their method to estimate cohesion values by using two relative abundance profiles of a training set and test set. Relative abundance profile of the training set was obtained by randomly selecting half of the samples in each microbiome passage. The test set consists of the other half of the samples. Using the training set and following the same procedure as Herren *et al*., connectedness metrics were calculated. The estimated connectedness metrics subtracts a null model. The objective of the null model was to calculate the strength of pairwise correlations that would be observed if there were no true relationship between OTUs. The obtained connectedness metrics are multiplied by relative abundance profile of test set to estimate positive and negative cohesion values. Two hundred iterations of sampling randomization in each microbiome passage were carried out at OTU level to obtain training set and test set for P1, P2, P3, and P4.

**Neutral model**

The neutral model was proposed by Sloan *et al*. to describe both microbial diversity and taxa-abundance distribution of a community (Sloan et al. 2007). Burns *et al*. (Burns et al. 2017) have developed a R package based on Sloan's neutral model to determine the potential importance of neutral process to a community assembly. In brief, the neutral model creates a potential neutral community by a single free parameter describing the migration rate, *m*, based on two sets of abundance profiles – a local community and metacommunities. The local community describes the observed relative abundance of OTUs, while the metacommunity is estimated by the mean relative abundance across all local communities. The estimated migration rate is the probability of OTU dispersal from the metacommunity to replace a randomly lost individual in the local community. The migration rate can be interpreted as dispersal limitation. In each microbiome passage, half of the samples were randomly selected and the relative abundance profile at the OTU level was used. The neutral model fit and migration rate were estimated in the resolution results of 200 iterations for P1, P2, P3, P4, and P4 Combined.

**Null model predictions**

We applied a null model approach on the serial passaging data P1-P4 to characterize the changes of stochastic process driving the assembly of plant microbiome over time. Lines that had high quality sequencing data at every time point (thirteen in total), were used for this analysis. The null scenario for each line at each passage was generated using the data for that same line at the previous passage. The null scenario of P1 was generated using the original field inoculum sample. The null model approach was based on community pairwise dissimilarity proposed by Chase and Myers (Chase and Myers 2011) and extended by Stegen *et al.* to incorporate species abundance (Stegen, Lin et al. 2013). Chase and Myers proposed a degree of species turnover by a randomization procedure where species probabilistically occur at each local community until observed local richness is reached. However, the estimated degree of turnover does not include species abundance. To take full advantage of our dataset, we also incorporated species relative abundance into the procedure proposed by Stegen *et al*.  Zinger *et al*. has developed R code for the null model and applied the null model approach on the soil microbiome (Zinger et al. 2019). This approach does not require *a priori* knowledge of the local community condition and determines if each plant microbiome at the current passage deviates from a null scenario generated by that same microbiome at the previous passage. In brief, the null scenario of each was generated by random resampling of OTUs and remained the same richness and number of reads with the original sample. Total OTUs observed in the sample and the corresponding relative abundance were used as probabilities of selecting an OTU and its associated number of reads, respectively. The Bray-Curtis distance is used to calculate dissimilarities across null communities with 1,000 permutations. The average of dissimilarities among permutations represents null expectations of community dissimilarities. The null deviation shows the differences between average null expectation and the observed microbiome of the same line.

Callahan, Benjamin J, Paul J McMurdie, Michael J Rosen, Andrew W Han, Amy Jo A Johnson, and Susan P Holmes. 2016. “DADA2: High-Resolution Sample Inference from Illumina Amplicon Data.” *Nature Methods* 13 (May): 581.

Carini, Paul, Patrick J. Marsden, Jonathan W. Leff, Emily E. Morgan, Michael S. Strickland, and Noah Fierer. 2017. “Relic DNA Is Abundant in Soil and Obscures Estimates of Soil Microbial Diversity.” *Nature Microbiology* 2 (3): 16242. https://doi.org/10.1038/nmicrobiol.2016.242.

Dixon, Philip, and M. W. Palmer. 2003. “VEGAN, a Package of R Functions for Community Ecology.” *Journal of Vegetation Science* 14 (6): 927–30. https://doi.org/10.1658/1100-9233(2003)014[0927:VAPORF]2.0.CO;2.

Edgar, Robert C., Brian J. Haas, Jose C. Clemente, Christopher Quince, and Rob Knight. 2011. “UCHIME Improves Sensitivity and Speed of Chimera Detection.” *Bioinformatics* 27 (16): 2194–2200. https://doi.org/10.1093/bioinformatics/btr381.

Herren, Cristina M., and Katherine D. McMahon. 2017. “Cohesion: A Method for Quantifying the Connectivity of Microbial Communities.” *The ISME Journal* 11 (11): 2426–38. https://doi.org/10.1038/ismej.2017.91.

Kozich, James J., Sarah L. Westcott, Nielson T. Baxter, Sarah K. Highlander, and Patrick D. Schloss. 2013. “Development of a Dual-Index Sequencing Strategy and Curation Pipeline for Analyzing Amplicon Sequence Data on the MiSeq Illumina Sequencing Platform.” *Applied and Environmental Microbiology* 79 (17): 5112–20. https://doi.org/10.1128/AEM.01043-13.

Lundberg, Derek S., Scott Yourstone, Piotr Mieczkowski, Corbin D. Jones, and Jeffery L. Dangl. 2013. “Practical Innovations for High-Throughput Amplicon Sequencing.” *Nature Methods* 10 (10): 999–1002. https://doi.org/10.1038/nmeth.2634.

McMurdie, Paul J., and Susan Holmes. 2013. “Phyloseq: An R Package for Reproducible Interactive Analysis and Graphics of Microbiome Census Data.” *PLOS ONE* 8 (4): e61217. https://doi.org/10.1371/journal.pone.0061217.

Morella, Norma M., Annika L. Gomez, Grant Wang, Michelle S. Leung, and Britt Koskella. 2018. “The Impact of Bacteriophages on Phyllosphere Bacterial Abundance and Composition.” *Molecular Ecology*, February. https://doi.org/10.1111/mec.14542.

Naylor, Dan, Stephanie DeGraaf, Elizabeth Purdom, and Devin Coleman-Derr. 2017. “Drought and Host Selection Influence Bacterial Community Dynamics in the Grass Root Microbiome.” *The ISME Journal* 11 (12): 2691. https://doi.org/10.1038/ismej.2017.118.

Nilsson, Rolf Henrik, Karl-Henrik Larsson, Andy F. S. Taylor, Johan Bengtsson-Palme, Thomas S. Jeppesen, Dmitry Schigel, Peter Kennedy, et al. 2019. “The UNITE Database for Molecular Identification of Fungi: Handling Dark Taxa and Parallel Taxonomic Classifications.” *Nucleic Acids Research* 47 (D1): D259–64. https://doi.org/10.1093/nar/gky1022.

Quast, Christian, Elmar Pruesse, Pelin Yilmaz, Jan Gerken, Timmy Schweer, Pablo Yarza, Jörg Peplies, and Frank Oliver Glöckner. 2013. “The SILVA Ribosomal RNA Gene Database Project: Improved Data Processing and Web-Based Tools.” *Nucleic Acids Research* 41 (D1): D590–96. https://doi.org/10.1093/nar/gks1219.

Rognes, Torbjørn, Tomáš Flouri, Ben Nichols, Christopher Quince, and Frédéric Mahé. 2016a. “VSEARCH: A Versatile Open Source Tool for Metagenomics.” e2409v1. PeerJ Preprints. https://doi.org/10.7287/peerj.preprints.2409v1.

. 2016b. “VSEARCH: A Versatile Open Source Tool for Metagenomics.” *PeerJ* 4 (October): e2584. https://doi.org/10.7717/peerj.2584.

Schloss, Patrick D., Sarah L. Westcott, Thomas Ryabin, Justine R. Hall, Martin Hartmann, Emily B. Hollister, Ryan A. Lesniewski, et al. 2009. “Introducing Mothur: Open-Source, Platform-Independent, Community-Supported Software for Describing and Comparing Microbial Communities.” *Applied and Environmental Microbiology* 75 (23): 7537–41. https://doi.org/10.1128/AEM.01541-09.

Skytte Andersen, Kasper S., Rasmus H. Kirkegaard, Soeren M. Karst, and Mads Albertsen. 2018. “Ampvis2: An R Package to Analyse and Visualise 16S RRNA Amplicon Data.” *BioRxiv*, January, 299537. https://doi.org/10.1101/299537.

Sloan, William T., Stephen Woodcock, Mary Lunn, Ian M. Head, and Thomas P. Curtis. 2007. “Modeling Taxa-Abundance Distributions in Microbial Communities Using Environmental Sequence Data.” *Microbial Ecology* 53 (3): 443–55. https://doi.org/10.1007/s00248-006-9141-x.

Takahashi, Shunsuke, Junko Tomita, Kaori Nishioka, Takayoshi Hisada, and Miyuki Nishijima. 2014. “Development of a Prokaryotic Universal Primer for Simultaneous Analysis of Bacteria and Archaea Using Next-Generation Sequencing.” *PLOS ONE* 9 (8): e105592. https://doi.org/10.1371/journal.pone.0105592.

Zinger, Lucie, Pierre Taberlet, Heidy Schimann, Aurélie Bonin, Frédéric Boyer, Marta De Barba, Philippe Gaucher, et al. 2019. “Body Size Determines Soil Community Assembly in a Tropical Forest.” *Molecular Ecology* 28 (3): 528–43. https://doi.org/10.1111/mec.14919.
